# Temporal gating of SIRT1 functions by O-GlcNAcylation prevents hyperglycemia and enables physiological transitions in liver

**DOI:** 10.1101/597153

**Authors:** Tandrika Chattopadhyay, Babukrishna Maniyadath, Hema P Bagul, Arindam Chakraborty, Namrata Shukla, Srikanth Budnar, Ullas Kolthur-Seetharam

## Abstract

Inefficient fasted-to-refed transitions are known to cause metabolic diseases. Thus, identifying mechanisms that may constitute molecular switches during such physiological transitions become crucial. Specifically, whether nutrients program a relay of interactions in master regulators, such as SIRT1, and affect their stability is underexplored. Here, we elucidate nutrient-dependent O-GlcNAcylation of SIRT1, within its N-terminal domain, as a key determinant of hepatic glucose- and fat-metabolism, and insulin signaling. SIRT1 glycosylation dictates interactions with PPARα/FOXO1/PGC1α/SREBP1, to exert a temporal control over transcription of genes during fasted-to-refed transitions. Interestingly, glycosylation-dependent cytosolic export of SIRT1 promotes a transient interaction with AKT and subsequent proteasomal degradation. Loss of glycosylation discomposes these interactions and enhances stability of SIRT1 even upon refeeding, which causes insulin resistance, hyperglycemia and hepatic-inflammation. Aberrant glycosylation of SIRT1 is associated with aging and/or metabolic diseases. Thus, nutrient-dependent glycosylation constrains spatio-temporal dynamics of SIRT1 and gates its functions to maintain metabolic homeostasis.

## Introduction

Metabolic states such as fed/fasted are a continuum and inability to mediate physiological transitions is often associated with diseases and aging (Chen et al., 1992; Dang et al., 2016; Geisler et al., 2016; Mezey, 1978; Petersen et al., 2017; Soeters et al., 2012). Most of our current understandings about molecular mechanisms that govern anabolic and catabolic processes have come from studies that investigated them as a comparative between fed and fasted states or as being distinct (Altarejos and Montminy, 2011; Chen et al., 1992; Horton et al., 1998; Naimi et al., 2010). However, given that these states are a continuum, unraveling nutrient/endocrine driven mechanisms that entail molecular switches and relay interactions becomes relevant. Specifically, it is important to investigate if/how changes in nutrient availability, which is aberrant during aging and in metabolic diseases, tunes molecular interactions to gate functions of master metabolic regulators during fed-fast-refed cycles.

Efficient metabolic transitions in the liver, which are majorly dependent on SIRT1, are vital for maintaining organism wide homeostasis and ensure constant supply of nutrients during fed-fast cycles (Banks et al., 2008; Dang et al., 2016; Geisler et al., 2016; Horton et al., 1998; Jitrapakdee, 2012; Pfluger et al., 2008; Purushotham et al., 2009; Rui, 2014; Xu et al., 2010). Abnormal activation or inhibition of hepatic fed- or fasting-responsive genes is known to cause metabolic diseases (Horton et al., 1998; Maniyadath et al., 2019; Mezey, 1978; Petersen et al., 2017). SIRT1, a fasting induced factor, is well established as a key regulator of liver functions that is largely mediated through deacetylation of transcription factors/co-activators (Ding et al., 2017; Houtkooper et al., 2012; Li, 2013; Yang et al., 2006). While deacetylation dependent activation of factors such as PGC1α, PPARα and FOXO1 is required for fatty acid oxidation and gluconeogenesis (Li, 2013; Sugden et al., 2010), inhibition of HIF1α and SREBP1 is known to repress expression of fed responsive genes (Lim et al., 2010; Ponugoti et al., 2010). Even though SIRT1 has been shown to be beneficial in countering aging and age-related diseases, intriguingly over-activation of SIRT1 functions is known to be detrimental (Kawashima et al., 2011; Vila et al., 2016; Wu et al., 2011). Hence, it is necessary to investigate mechanisms that control SIRT1 turnover and homeostasis under normal and pathophysiological states.

It is important to note that SIRT1, although being induced during fasting, is necessary to activate insulin signaling by regulating almost all the components of the pathway and a loss of SIRT1 leads to insulin resistance (Liang et al., 2009; Wang et al., 2011). Specifically, SIRT1 is known to deacetylate and activate AKT, the nodal kinase in insulin signaling (Sundaresan et al., 2011). Importantly, as SIRT1 is largely nuclear, spatio-temporal regulation of SIRT1-AKT interactions during such state transitions is yet to be unraveled. Moreover, paradoxically, SIRT1 seems to counter the induction of lipogenic and glycolytic genes, which are downstream to insulin signaling (Ding et al., 2017; Ponugoti et al., 2010; Rodgers and Puigserver, 2007). Despite these, it is still unclear if/how interactions of SIRT1 with both transcriptional regulators and signaling factors are temporally resolved during fed-fast cycles. Importantly, the mechanism that operates to either engage or disengage SIRT1 from its substrates has not been elucidated, yet.

While SIRT1 is known to interact with a plethora of nuclear and cytosolic factors, it is only recently that mechanisms, which may determine specificity of interactions, have begun to be unraveled. In this regard, others and we have recently highlighted the importance of the N-terminal domain of SIRT1 (Deota et al., 2017; Ghisays et al., 2015; Krzysiak et al., 2018; Pan et al., 2012). Notably, we have found that a domain encoded by exon-2 in SIRT1, which forms intrinsically disordered region (IDR), acts to tether substrate proteins and is specifically required for interaction with PGC1α, PPARα and FOXO1 (Deota et al., 2017). Given that modifications on IDRs are known to determine protein-protein interactions (Babu, 2016; Oldfield and Dunker, 2014), it is enticing to hypothesize SIRT1-IDR modification as a means to regulate interactions, especially during metabolic transitions.

Dynamic and reversible O-GlcNAcylation (glycosylation) on serine/threonine residues, has emerged as a major regulator of nuclear and cytosolic proteins involved in diverse cellular functions (Bond and Hanover, 2013; Hart, 1997; Wells et al., 2003). The donor metabolite for glycosylation, UDP-GlcNAc, is derived from hexosamine biosynthetic pathway along with contributions from various anabolic pathways and is therefore considered to act as a signal of a nutrient replete state (Chiaradonna et al., 2018; Zachara and Hart, 2006). O-GlcNAc-transferase (OGT) has been identified as the only enzyme that mediates glycosylation across cellular compartments and O-GlcNAcase (OGA) mediates deglycosylation (Butkinaree et al., 2010; Slawson et al., 2006). Further, OGT dependent glycosylation is enhanced following insulin stimulation and glucose uptake (Whelan et al., 2008).

In the present study, we show that SIRT1 is modified under nutrient replete conditions via O-GlcNAcylation and aberrant SIRT1 glycosylation could potentially be a major factor contributing to metabolic diseases and aging. Interestingly, upon refeeding nutrient dependent glycosylation occurs within a domain that encodes substrate specificity and thus, switches its interaction from transcriptional factors/co-regulators to the nodal kinase in insulin signaling. Molecular and physiological assays clearly show that glycosylation of SIRT1 is critical to spatio-temporally resolve its nuclear and cytosolic functions, which is required for efficient fed-fast-refed transitions. Furthermore, glycosylation also causes SIRT1 degradation via ubiquitin-proteasomal pathway and limits its function in a fed state. Besides identifying a molecular mechanism that constrains SIRT1 functions, we establish that active glycosylation-de-glycosylation cycles in SIRT1 are essential for liver functions and maintenance of metabolic homeostasis.

## Results

### Spatial and temporal shift in SIRT1 interactions mediated by nutrient inputs

We sought out to address if metabolic inputs orchestrate SIRT1 interactions and functions in response to changes in nutrient availability, such as during fast-to-refed transitions. Towards this, we examined interaction of SIRT1 with transcription factors like FOXO1 and PPARα, and co-activators such as PGC1α, which are essential to mediate a fasting response. As shown in Figure 1A, SIRT1 interaction with these factors reduced notably under high glucose conditions when compared to cells grown in low glucose medium. Interestingly, we also found a similar decrease in interactions with SREBP1, a well-known mediator of a fed response (Figure 1B). Among others, interaction of SIRT1 with AKT, a nodal kinase downstream to insulin, is known to activate insulin signaling. Moreover, loss of SIRT1 has been shown to induce insulin resistance (Liang et al., 2009; Schenk et al., 2011; Sundaresan et al., 2011; Wang et al., 2011). Therefore, we wondered if glucose and/or insulin affected the ability of SIRT1 to interact with AKT. It was very interesting to see that addition of insulin to cells led to a temporal regulation of SIRT1-AKT interactions with a significant increase at 15 minutes, which thereafter decreased drastically by 30 minutes (Figure 1C). These indicated that while fed inputs (nutrient/endocrine) led to a disruption of SIRT1 interactions with nuclear transcription factors and co-activators, there was a transient increase in binding with a cytosolic kinase, AKT.

**Figure 1:**
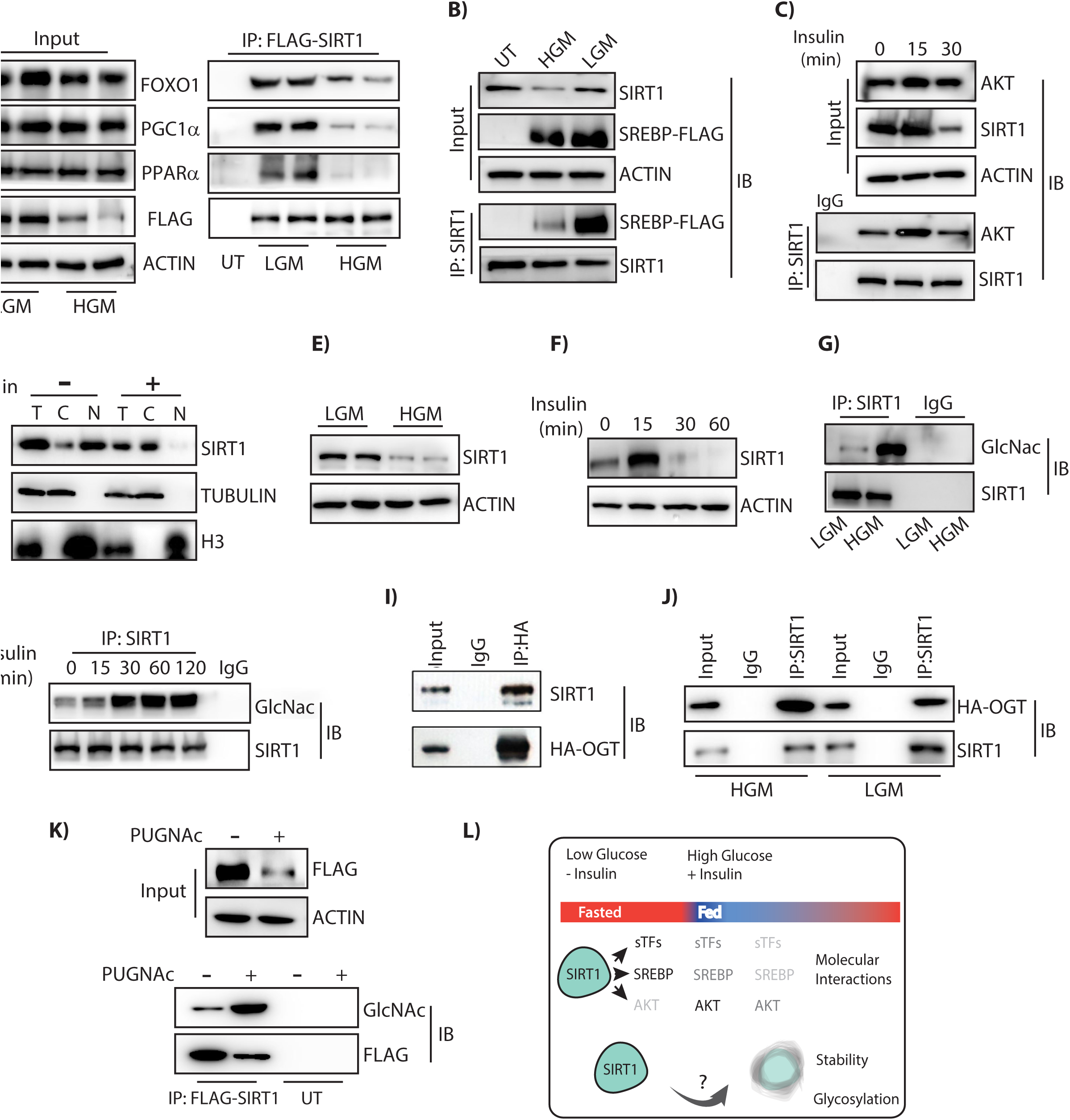
Nutrient dependent differential interaction, stability and glycosylation of SIRT1: (A-C) Immunoblot for interactions in co-immunoprecipitation (A) using FLAG-M2 beads from HEK293T cells expressing FLAG-SIRT1-WT in response to 12 h of low glucose (LG) and high glucose (HG) culture conditions; (B) of SIRT1 from HEK293T cells expressing FLAG-SREBP1 in response to 12 h of LG and HG culture conditions; (C) of SIRT1 from HEK293T cells treated with 100nM Insulin for indicated time points. (D-F) Immunoblot for endogenous SIRT1 levels in (D-E) Primary hepatocytes treated with (D) Insulin (100nM) and (E) LG and HG media for 12 h; (F) Mouse liver tissue, harvested at indicated time points in response to Insulin (0.75 IU/kg b.wt) administered intraperitoneally. (G-H) Immunoblot for glycosylated SIRT1 in (G) Primary hepatocytes treated with 12h of LG and HG media and (H) Mouse liver tissue harvested at indicated time points upon Insulin (0.75 IU/kg b.wt) administration. (I-J) Immunoblot for endogenous SIRT1 in HA immunoprecipitates from HEK293T cells expressing (I) HA-OGT and (J) HA-OGT treated with LG and HG media. (K) Immunoblot for glycosylated SIRT1 in FLAG- M2 immunoprecipitates from HEK293T cells expressing FLAG-SIRT1-WT treated with 100µM PUGNAc for 16 h. (L) Schematic representation of differential SIRT1 interactions and stability under fasted and fed state. T=Total cell lysate; C=Cytoplasmic fraction; N= Nuclear fraction. *See also Figure S1*.

These results prompted us to investigate if the localization of SIRT1 is affected under fed and starved conditions. Although, SIRT1 has been shown to shuttle between nucleus and the cytoplasm (Nasrin et al., 2009; Tanno et al., 2007; Tong et al., 2013), a nutrient dependent control of SIRT1 localization is largely unknown. Insulin administration under high glucose resulted in re-localization of SIRT1 from nucleus to the cytoplasm (Figure 1D). Incidentally, SIRT1 levels not only decreased following a normal physiological transition to a fed state (Figures 1A-1F and S1A-S1C) but also in livers of high fat diet (HFD) and aged mice (Figures S1D and S1E). Although, SIRT1 levels are known to decrease during a fed state (Kanfi et al., 2008; Rodgers et al., 2005), if/how nutrient inputs regulate SIRT1 homeostasis is still unknown. Together, our results pointed towards a glucose/nutrient dependent control of SIRT1 dynamics in terms of interactions, localizations and expression or stability.

### SIRT1 is glycosylated during a fed state

We then hypothesized that a nutrient derived post translational modification that is sensitive to cellular levels of glucose could potentially regulate SIRT1. O-GlcNAcylation or glycosylation has been proposed to link nutrient excess states to molecular functions of a diverse set of intracellular proteins. Thus, we investigated if altered SIRT1 dynamics in starved and fed states could be mediated by glycosylation. Interestingly, we found SIRT1 to undergo glycosylation in response to both insulin administration and under high-glucose conditions (Figures 1G and 1H). We confirmed SIRT1 glycosylation by reciprocal pull-downs viz. assaying for glycosylation on immunoprecipitated SIRT1 and by checking for SIRT1 enrichment on sWGA-beads (which pulls down all O-GlcNAcylated proteins) (Figures 1G, 1H and S1F). Importantly, glycosylated SIRT1 levels diminished when insulin stimulation was done in the presence of 2-Deoxy-Glucose (2-DG), a competitive inhibitor of glycolysis (Figure S1G). These results clearly indicate that glucose metabolism is essential for glycosylation of SIRT1 under fed conditions.

OGT catalyzes the addition of O-linked beta-N-acetylglucosamine (Butkinaree et al., 2010). Enhanced glycosylation of SIRT1 under a glucose replete condition prompted us to probe for OGT-SIRT1 interactions under altered metabolic states. We found that SIRT1 does interact with OGT (Figure 1I), and interestingly, this was substantially higher under fed conditions (Figure 1J) and decreased in the presence of 2-DG (Figure S1H). These results suggested that a fed response led to glucose dependent enhanced glycosylation of SIRT1 mediated by OGT. Inhibiting either the deglycosylase enzyme, OGA (using PUGNAc) or OGT (using BZX) led to a significant increase and decrease in SIRT1 glycosylation, respectively, implying this to be an active process (Figures 1K and S1I).

Together, these findings indicated that SIRT1 interactions, localization, stability and glycosylation were correlated with a metabolic shift. Therefore, we surmised that a nutrient dependent modification of SIRT1 could be orchestrating its dynamics at a molecular level, which enables physiological transitions and thus play an important role in maintaining homeostasis (Figure 1L). We then set out to test this systematically, as described below.

### SIRT1 is glycosylated within its specificity domain

In silico prediction of potential glycosylation sites within SIRT1 indicated that Thr^160^/Ser^161^ (T^160^/S^161^) within the exon-2 domain could be modified (Figure S2A). Importantly, this domain, which also happens to be intrinsically disordered, has been identified as a key determinant of binding specificities of SIRT1 with transcription factors (Deota et al., 2017; Krzysiak et al., 2018). We therefore predicted that glycosylation at T^160^/S^161^ in mouse SIRT1 (mSIRT1) (Figure 2A) could play a critical role in regulating SIRT1 functions. Importantly, this domain is highly conserved across mammals with 84% identity between humans and mice, and the predicted glycosylation site is conserved across species (Ser^161^/Ser^169^ in mouse/human, respectively) (Figure 2A). Thus, we mutated both T^160^/S^161^of mSIRT1 to Alanine, i.e. T^160^/S^161^-A, henceforth termed as mutant-E2 (mutE2) (Figure S2B) and found its expression to be unaltered under low and high glucose conditions (Figure 2B). Reversible pull-down with FLAG-M2 and sWGA beads clearly showed that glycosylation of SIRT1- T^160^/S^161^-A (SIRT1-mutE2) was starkly reduced (Figures 2C and 2D). Interestingly, we found that the SIRT1 exon-2 domain was necessary to mediate the interaction with OGT and in the absence of exon-2 domain SIRT1 glycosylation was markedly reduced (Figures S2C and S2D). However, it is important to note that the decreased glycosylation in SIRT1-mutE2 was not due to reduced interaction with OGT but a resultant of loss of glycosylatable residues T^160^/S^161^ within the exon-2 domain (Figure S2C). Based on our predictions and a concurrent study (Han et al., 2017), we surmise that other sites on SIRT1 could contribute to the residual glycosylation on SIRT1-mutE2.

**Figure 2:**
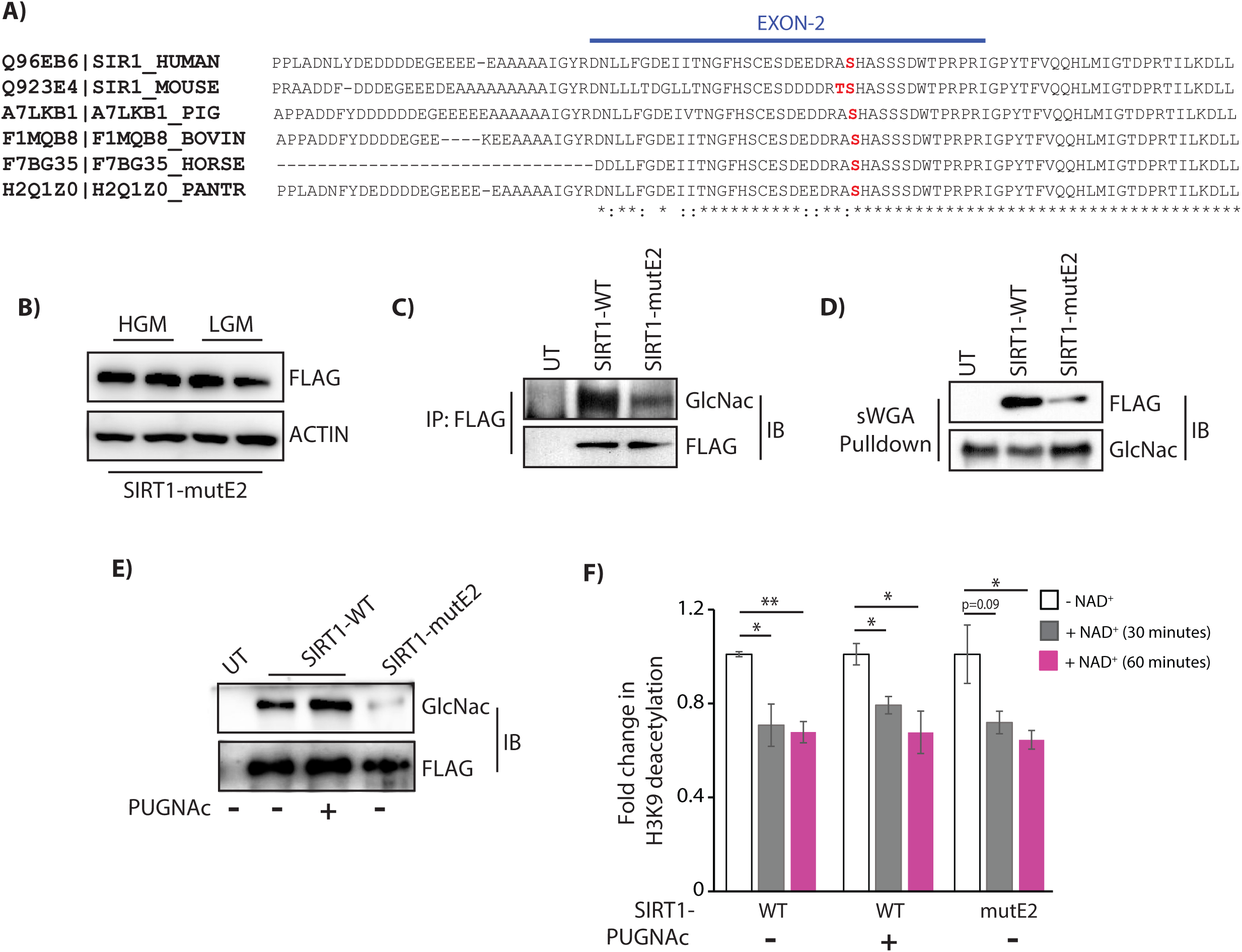
Identification of SIRT1 glycosylation sites in Exon-2 region of SIRT1: (A) ClustalW alignment of amino acid sequence of SIRT1-WT from human, mouse, pig, bovine, horse and panther respectively. Serine at position 161 of SIRT1 in the Exon 2 region, which is evolutionarily conserved, has been indicated in red. (B) Immunoblot for FLAG-SIRT1-mutE2 in HEK293T cells grown in LG and HG culture media. (C-D) Immunoblot for glycosylated SIRT1 from HEK293T cells expressing FLAG-SIRT1-WT and FLAG-SIRT1-mutE2 (C) in FLAG-M2 immunoprecipitates and (D) sWGA pull down. (E-F) Comparison of SIRT1 activity on acetylated histones along with saturating levels of NAD^+^ for 30 and 60 minutes from FLAG-M2 immunoprecipitates in HEK293T cells overexpressing FLAG-SIRT1-WT (± PUGNAc) and – mutE2 (E) Immunoblotted for glycosylated SIRT1 and (F) Bar graph for acetylated H3K9 levels normalized to FLAG (n=3). Data are represented as mean ± SEM and analyzed by the Student’s t test. A value of P ≤ 0.05 was considered statistically significant. *P≤ 0.05; **P≤ 0.01; ***P≤ 0.001. *See also Figure S2*.

Post-translational modifications on SIRT1 such as phosphorylation, sumoylation, etc. are known to modulate its activity (Ford et al., 2008; Gao et al., 2011; Peng et al., 2015; Revollo and Li, 2013). Hence, to investigate whether glycosylation resulted in any difference in SIRT1 activity, we carried out catalytic assays using immunoprecipitated SIRT1-WT and SIRT1-mutE2, as indicated. On comparing NAD^+^-dependent SIRT1 mediated loss of lysine-9 acetylation of histone H3 (H3K9Ac), *in vitro*, we found comparable activities for both these forms of SIRT1 (Figures 2E-2F and S2E-S2F). This was not surprising given that we had earlier found that loss of exon-2 domain, which although was necessary for modification of endogenous proteins by differential tethering, did not alter catalytic activity of SIRT1 on accessible acetylated peptides (Deota et al., 2017). These suggested that glycosylation or a loss of it did not significantly impact the intrinsic activity on peptide substrates. However, given that it is nearly impossible to obtain SIRT1, which is stoichiometrically enriched in glycosylation at only these sites, we cannot rule out possible effects of this modification on SIRT1 activity. Nevertheless, while these results clearly established that T^160^/S^161^ within the exon-2 domain were the predominant nutrient sensitive glycosylation sites, they also raised the possibility of this modification regulating SIRT1 binding with its interacting proteins in fed-fast conditions.

### SIRT1 glycosylation determines molecular interactions with nuclear and cytosolic factors

SIRT1 interacts with and deacetylates a multitude of transcription factors and thus exerts a master control over gene expression (Li, 2013; Rodgers and Puigserver, 2007; Sugden et al., 2010). Hence, to check if glycosylation was involved in integrating nutrient inputs with SIRT1 functions, we assayed for the ability of wild type and mutE2 forms of SIRT1 to interact with transcription factors and co-activators in cells that were treated with/without PUGNAc. Surprisingly, we found that glycosylation on SIRT1 notably reduced its interaction with transcriptional regulators that operate during, starved and fed conditions (Figures 3A and 3B). On the contrary, the non-glycosylatable form of SIRT1 (i.e., mutE2) had enhanced interactions with these factors (Figures 3C and 3D). These findings indicate that hypo-glycosylation of SIRT1, at T^160^/S^161^ under starved or nutrient deprived conditions, increases its association with PGC1α, FOXO1, PPARα and SREBP1, to possibly up regulate the expression of genes required for starvation response and simultaneously suppresses the expression of fed responsive genes.

**Figure 3:**
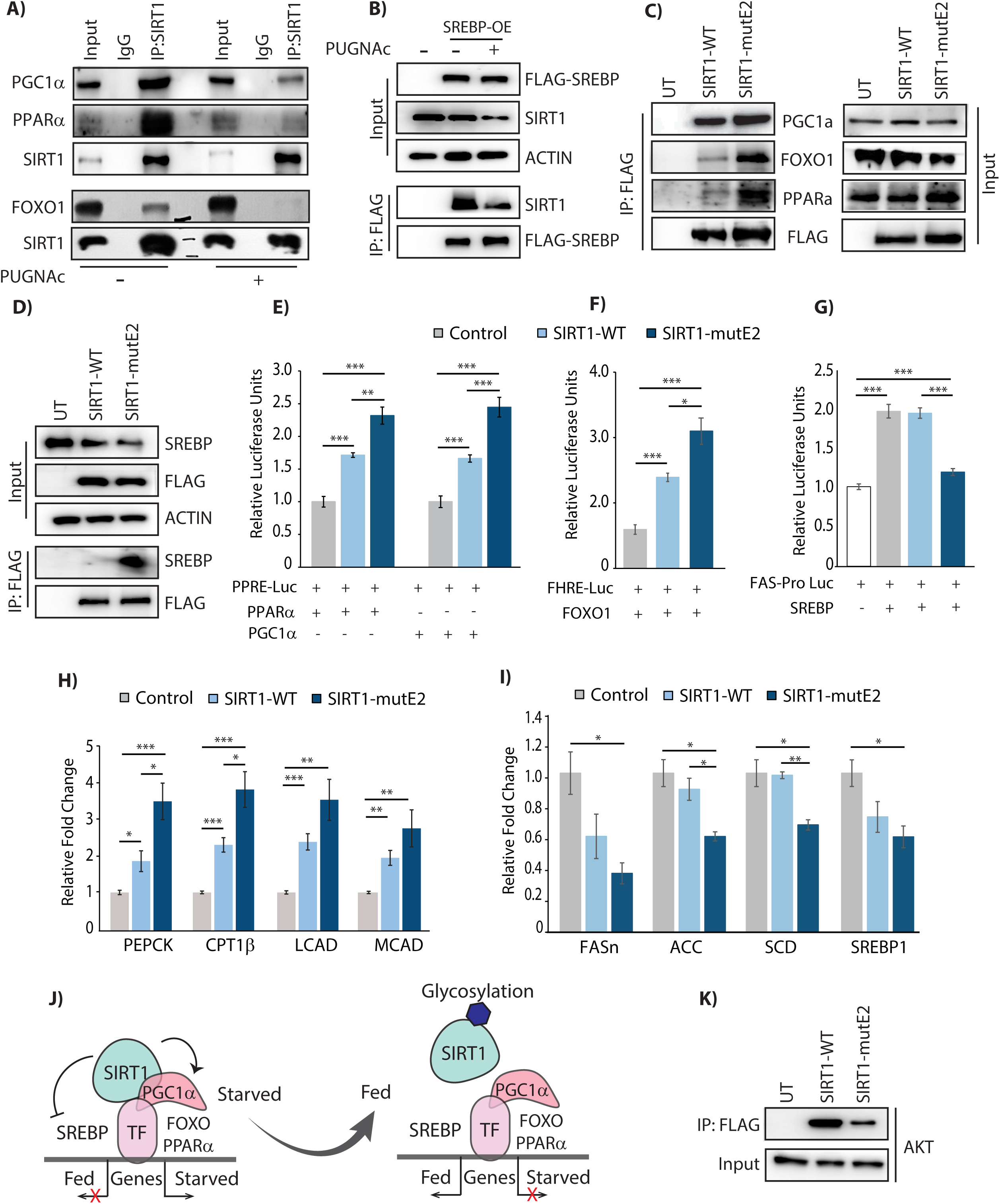
SIRT1 glycosylation modulates interactions and affects transcription: (A-D) Immunoblot for interactions in (A-B) HEK293T cells treated with 100µM PUGNAc for 12-16 h from (A) SIRT1 immunoprecipitates and (B) FLAG-M2 immunoprecipitates overexpressing FLAG-SREBP1; (C-D) Immunoblot of FLAG-M2 immunoprecipitates from HEK293T cells expressing either FLAG-SIRT1-WT or -mutE2. (E-G) Luciferase assay in HEK293T cells expressing either FLAG-SIRT1-WT or -mutE2 and quantifying (E) PGC1α or PPARα - dependent transcription as measured by PPRE-luciferase assay (n=3-4, N=3); (F) FOXO1 dependent transcription as measured by FHRE-Promoter-luciferase assay (n=3, N=2) and (G) SREBP1 dependent transcription as measured by FAS-Promoter-luciferase assay (n=3-4, N=2). (H-I) Gene expression analysis from HepG2 cells expressing either FLAG-SIRT1-WT or -mutE2 (n=3, N=2) (H) Gluconeogenic and beta-oxidation genes and (I) Lipogenic genes. (J) Schematic representation of effect of glycosylation on SIRT1 dependent molecular alterations in fasted and fed state. (K) Immunoblot for AKT interaction in FLAG-M2 immunoprecipitates from HEK293T cells expressing either FLAG-SIRT1-WT or -mutE2. Data are represented as mean ± SEM and analyzed by the Student’s t test. A value of P ≤ 0.05 was considered statistically significant. *P≤ 0.05; **P≤ 0.01; ***P≤ 0.001. *See also Figure S3*.

To ascertain this, we assayed for the expression of genes downstream to these transcription factors, using both luciferase based reporter constructs and by measuring endogenous mRNA levels of fasting-/fed-responsive genes. We found that non-glycosylated SIRT1 enhanced transcription downstream to PGC1α, PPARα and FOXO1, including genes involved in oxidative phosphorylation, gluconeogenesis and beta-oxidation, and inhibited expression of lipogenic genes downstream to SREBP1 (Figures 3E-3I). Furthermore, increased palmitate induced fatty acid oxidation and ATP production rate was observed in hepatocytes expressing SIRT1-mutE2 compared to SIRT1-WT (Figures S3A and S3B). In addition to corroborating the differential interactions of unmodified and modified SIRT1, these results demonstrate that nutrient dependent SIRT1 glycosylation is necessary to switch the transcription program, for efficient transition from a starved to a fed state (Figure 3J).

SIRT1 is well documented to be a key factor that determines insulin sensitivity (Liang et al., 2009; Ramachandran et al., 2011). In fact, SIRT1 mediated deacetylation of AKT is important for its membrane localization and activity in response to insulin stimulation (Sundaresan et al., 2011). However, spatial and temporal regulation of SIRT1-AKT interactions during starved to refed transition is still unclear. In this context, we found that SIRT1 interaction with AKT was enhanced at initial phases post insulin and glucose administration, and coincided with SIRT1 glycosylation (Figure 1C). To check if glycosylation was necessary for SIRT1-AKT interactions, we assayed this using SIRT1-mutE2. It is evident from Figure 3K, that in the absence of glycosylation at T^160^/S^161^, SIRT1 failed to bind to AKT. In addition, overexpression of this non-glycosylatable form of SIRT1 resulted in dampened insulin signaling when compared to SIRT1-WT (Figures S3C), corroborating the differential interaction observed in Figure 3K. Interestingly, cells expressing SIRT1-mutE2 also displayed reduced glycolytic flux under both basal and insulin stimulated conditions, clearly highlighting the metabolic consequence of reduced insulin signaling upon loss of glycosylation at T^160^/S^161^ (Figures S3D-S3E). This illustrates that nutrient driven modification of SIRT1, a fasting factor, during the initial phase of a refed response is required for activation of insulin signaling.

Moreover, it was interesting to note that SIRT1-AKT interactions decreased during the latter phase of a refed response (Figure 1C). Together with reduced SREBP1 interactions, this indicates that while nutrient dependent glycosylation in SIRT1 is required for insulin signaling, it also ensures that a fed responsive transcriptional program is not inhibited by SIRT1. This is a significant finding since it provides molecular basis to the paradoxical role of SIRT1 in activating insulin signaling and inhibiting the downstream fed transcriptional program.

### Glycosylation results in nuclear export of SIRT1

The dynamic switch in loss and gain of SIRT1 interactions with nuclear transcription factors and cytoplasmic signaling molecules, respectively, in response to insulin/glucose administration, was striking. This prompted us to ask if glycosylation affected nuclear-cytoplasmic distribution of SIRT1. Consistent with earlier reports, we found that SIRT1 was predominantly nuclear (Figure 4A). As anticipated, SIRT1 re-localized to cytoplasm in PUGNAc treated cells under high glucose, and negligible protein was seen in the nucleus (Figure 4A). Further, glycosylated SIRT1 was enriched in the cytoplasmic fraction of sWGA pull downs (Figure S4A). Moreover, we saw enhanced nuclear retention of SIRT1-WT upon inhibition of glycosylation even under nutrient excess conditions (Figure 4B) and there was stark contrast between localization patterns of SIRT1-WT and SIRT1-mutE2 (Figure 4C), with predominant nuclear localization in the absence of glycosylation at T^160^/S^161^. These results clearly established that nutrient dependent glycosylation determined nuclear-cytosolic pools of SIRT1.

**Figure 4:**
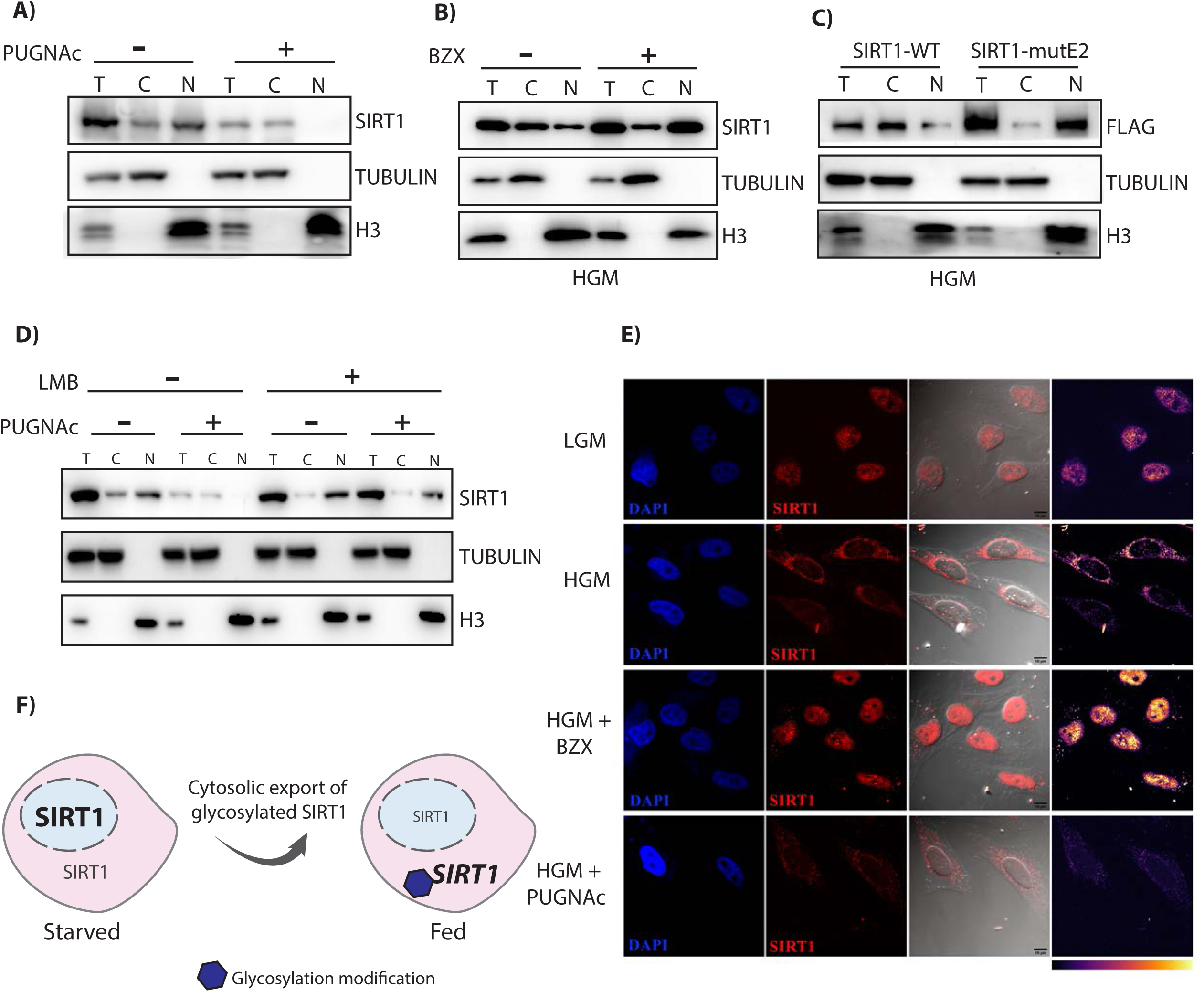
N-terminal glycosylation dependent cytoplasmic export of SIRT1: (A-D) Nucleo-Cytoplasmic distribution of SIRT1 in HeLa cells (A) treated with 100µM PUGNAc for 12 h and (B) treated with 100µM BZX for 12 h under HG conditions; (C) expressing either FLAG-SIRT1- WT or -mutE2 under HG conditions and (D) Treated with 100µM PUGNAc and 10ng/ml LMB for 12 h. (E) Representative immunofluorescence image for endogenous SIRT1 localization upon treatment (16 h) with either OGT inhibitor (BZX) or OGA inhibitor (PUGNAc) in HeLa cells grown in LG or HG media, as indicated (n=3-4). (F) Schematic representation of cytosolic export of SIRT1 upon nutrient dependent glycosylation at the N-terminus. T=Total cell lysate; C=Cytoplasmic fraction; N= Nuclear fraction. *See also Figure S4*.

Next, we addressed if SIRT1 was glycosylated within the nucleus and if the relocalization was an active mechanism. Cells treated with Leptomycin B (LMB), a known inhibitor of exportin protein, CRM-1, resulted in loss of nuclear extrusion of SIRT1, which indicated that it was an active process under a fed state (Figures 4D and S4B). OGT is known to localize to the nucleus, apart from the cytoplasm and mitochondria, and various nuclear proteins are known to be glycosylated (Butkinaree et al., 2010; Issad and Kuo, 2008; Yang and Qian, 2017). On assaying for glycosylation of SIRT1 immunoprecipitated from LMB treated cells under high glucose, we found that SIRT1 was indeed modified within the nuclear compartment (Figure S4C). Thus, we establish that nutrient dependent glycosylation of SIRT1 at its N-terminal domain results in its cytoplasmic export (Figures 4E and 4F).

### Nutrient dependent degradation of SIRT1 is mediated by glycosylation

While mechanisms that induce SIRT1 during a fasting phase are well known (Kwon and Ott, 2008; Revollo and Li, 2013), relatively little is known about those that lead to its degradation during a fed state. Importantly, if/how nutrient status of the cells control SIRT1 turnover remains to be unraveled. Treatment of cells with the translational inhibitor cyclohexamide (CHX) resulted in increased degradation of SIRT1 in the presence of PUGNAc, when compared to untreated cells (Figure 5A) and suggested that glycosylation decreased SIRT1 stability. In addition, stability of SIRT1 was markedly reduced upon CHX treatment in the presence of insulin (Figure 5B). Interestingly, while inhibiting OGA led to enhanced degradation of SIRT1, inhibiting OGT (using BZX) that mediates glycosylation stabilized SIRT1 even under fed conditions (Figure S5A). Together, these results clearly illustrate that nutrient dependent glycosylation was involved in regulating SIRT1 levels. To further identify whether SIRT1 degradation was proteasome mediated, cells were treated with proteasomal inhibitor MG132 in the presence or absence of insulin. As shown in Figure 5C, SIRT1 degradation was abrogated in the presence of MG132. Immunoblotting for ubiquitination of immunoprecipitated SIRT1 under these conditions clearly showed that sustained glycosylation resulted in increased ubiquitination in cells treated with PUGNAc and MG132 (Figure 5D). However, it has been difficult to tease out the temporal switch from glycosylation to ubiquitination during the continued fed condition. Moreover, whether nutrient dependent inputs activate specific E3-ligases needs to be investigated in the future. Importantly, we found that the stability of SIRT1-mutE2 was significantly enhanced as compared to the wild type protein, even in the presence of PUGNAc (Figure S5B). The stability of SIRT1-mutE2 protein was similar between PUGNAc treated and untreated cells indicating that the observed effect on protein turnover was primarily driven by glycosylation at T^160^/S^161^ (Figure 5E). Interestingly, nuclear retention of SIRT1-WT, in LMB and PUGNAc treated cells, did not lead to increased destabilization and suggested that cytoplasmic export was necessary for its degradation (Figures S5C and S5D). Together, these conclusively demonstrate that glycosylation leads to ubiquitin mediated degradation of SIRT1 under nutrient rich conditions, which was hitherto unknown (Figure 5F).

**Figure 5:**
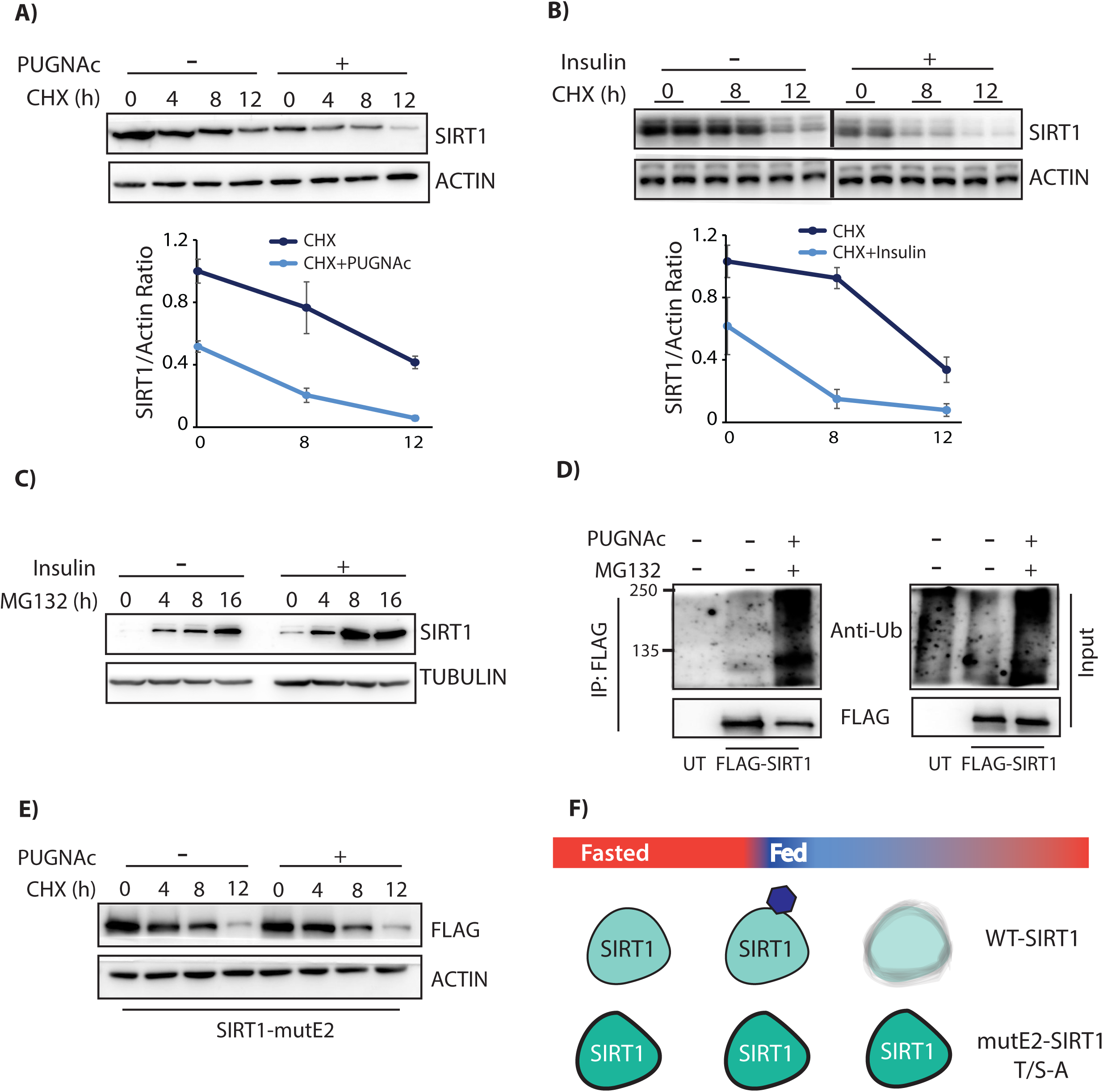
Glycosylation affects SIRT1 stability: (A-C) Immunoblot for endogenous SIRT1 levels in HepG2 cells treated with (A) PUGNAc (100µM, 16 h) and Cyclohexamide (CHX, 100µg/ml) for indicated time points (n=4); (B) Insulin (100nM, 16 h) and Cyclohexamide (CHX, 100µg/ml) for indicated time points (n=4) and (C) Insulin (100nM, 16 h) and proteasomal inhibitor MG132 (20µM) for indicated time points. (D) Immunoblot of ubiquitinated SIRT1 from SIRT1 immunoprecipitates in HepG2 cells treated with PUGNAc and MG132 for 16 h. (E) Immunoblot for SIRT1 levels in HepG2 cells expressing either FLAG SIRT1-WT or -mutE2 and treated with PUGNAc (100µM, 16 h) and Cyclohexamide (CHX, 100µg/ml) for indicated time points. (F) Schematic representation of SIRT-WT and SIRT-mutE2 protein stability under fasted and fed states. Data are represented as mean ± SEM and analyzed by the Student’s t test. A value of P ≤ 0.05 was considered statistically significant. *P≤ 0.05; **P≤ 0.01; ***P≤ 0.001. *See also Figure S5*.

### Reduced SIRT1 expression during aging and obesity is associated with glycosylation

Reduced SIRT1 expression has been associated with aging and metabolic diseases, both causally and consequentially. While decreased transcription has been reported (Kwon and Ott, 2008; Longo and Kennedy, 2006; Zschoernig and Mahlknecht, 2008), whether a nutrient dependent input/modification results in reduced protein stability and thus, lower expression under these conditions has not been elucidated, yet. On assaying for SIRT1 expression we found that there was a drastic reduction in young refed, high-fat-induced obese and old mice livers, consistent with our earlier results and previous reports (Figures S1C-S1E) (Gao et al., 2011; Kanfi et al., 2008; Maniyadath et al., 2019). Notably, on loading equal amounts of immunoprecipitated SIRT1 from these mice livers, we found a significant increase in SIRT1 glycosylation, as compared to their respective controls (Figures 6A-6C). Although correlative, these suggest that glycosylation of SIRT1 could possibly be one of the underlying causes for its reduced expression during aging or in obesity.

**Figure 6:**
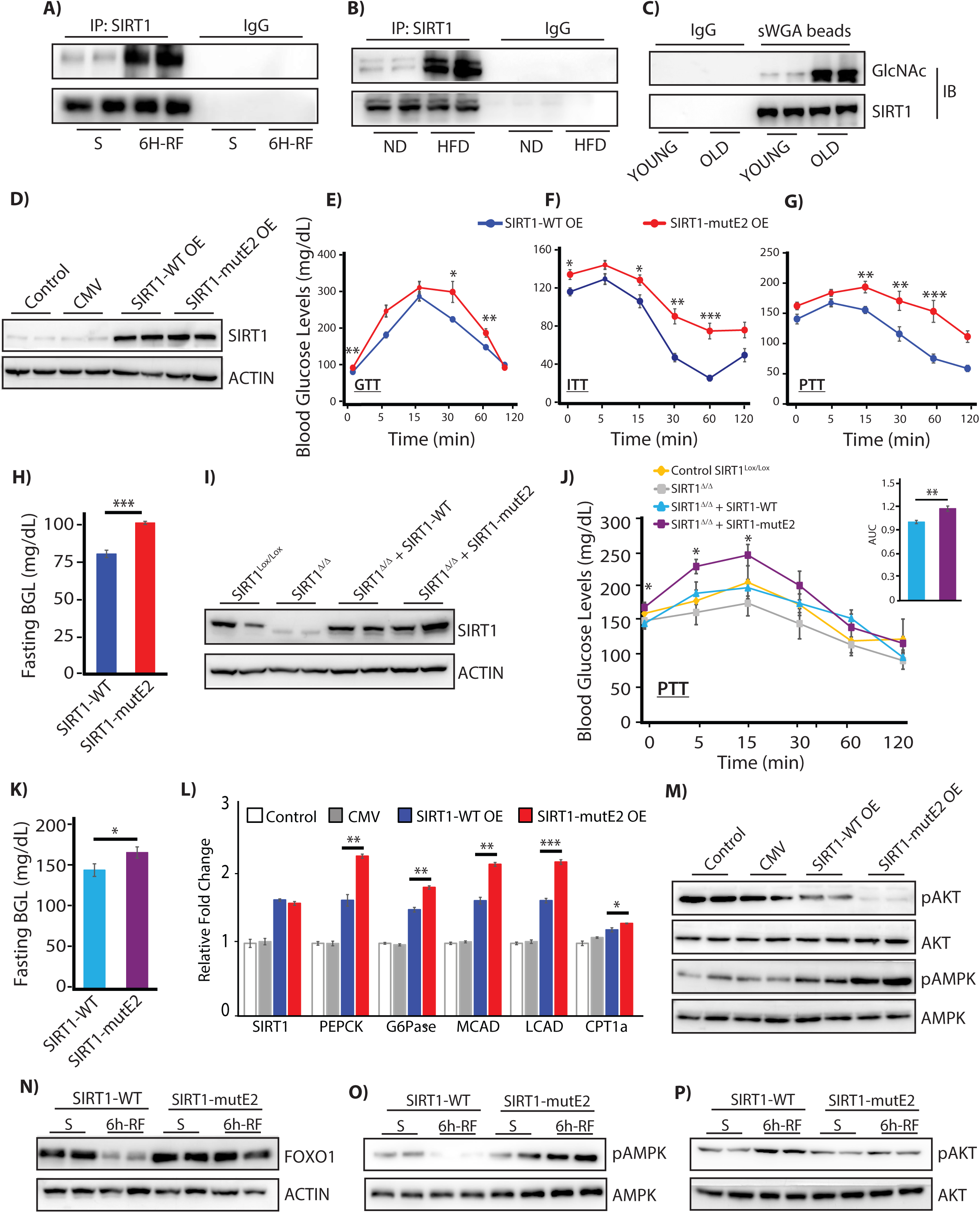
SIRT1 glycosylation is essential for liver homeostasis and starved-refed transition: (A-C) Immunoblot of glycosylated SIRT1 from SIRT1 immunoprecipitates from livers of (A) 24 h starved and 6 h refed; (B) Normal fed chow diet (ND) and 3 months fed High Fat Diet (HFD) and (C) Young (3 months) and old (20 months) mice. (D-H) Physiological assays following overexpressing of either SIRT1-WT or SIRT1-mutE2, along with control injected and uninjected mice (D) Immunoblot of SIRT1 in the liver; (E) Glucose tolerance test (GTT), (F) Insulin tolerance test (ITT) and (G) pyruvate tolerance test (PTT) (n=6, N=3); (H) Blood glucose levels upon 12 h of fasting (n=6, N=3). (I-K) Physiological assays in SIRT1^co/co^ mice rescued with either SIRT1-WT or SIRT1-mutE2 after floxing out endogenous SIRT1 along with uninjected control (I) Immunoblot for SIRT1 in liver tissues; (J) Pyruvate tolerance test and area under the curve (AUC) (n=6) and (K) Blood glucose levels upon 12 h of fasting (n=6); (L-M) Liver tissues harvested from mice with hepatic overexpression of either SIRT1-WT or SIRT1-mutE2, along with relevant controls, and analyzed for (L) Gluconeogenic and beta-oxidation gene expression (n=3, N=2) and (M) Immunoblot of pAMPK and pAKT for fasted and fed signaling, with total AMPK/AKT. (N-P) Immunoblot of 24 h starved and 6 h-refed liver tissues harvested from SIRT1^co/co^ mice rescued with either SIRT1-WT or SIRT1-mutE2 after floxing out endogenous SIRT1, as indicated. Data are represented as mean ± SEM and analyzed by the Student’s t test. A value of P ≤ 0.05 was considered statistically significant. *P≤ 0.05; **P≤ 0.01; ***P≤ 0.001. *See also Figure S6*.

### Non-glycosylated SIRT1 leads to hyperglycemia and abrogates fasted-to-refed transition

To unravel the physiological significance of SIRT1 glycosylation, we overexpressed both wild type and SIRT1-mutE2 in livers of C57Bl6 mice using adenovirus (Figures S6A and 6D). We found that mice with ectopic expression of SIRT1-mutE2 led to disrupted glucose homeostasis as evident from glucose-, insulin- and pyruvate-tolerance tests (GTT, ITT and PTT respectively) (Figures 6E-6G). Further, we found a significant increase in fasting blood glucose in mice overexpressing SIRT1-mutE2 (Figure 6H), and together with the PTT results this clearly indicated increased gluconeogenesis. In order to negate the possible artifacts of overexpression, we knocked out endogenous SIRT1 in the livers of SIRT1^lox/lox^ mice and restored it with either wild type or mutE2 forms using adenoviral systems (Figure S6B). As shown in Figure 6I, both the SIRT1 forms were expressed to amounts that were comparable to the endogenous SIRT1 in the liver. It is important to note that similar to ectopic overexpression, SIRT1 floxed out mice, expressing SIRT1-mutE2 had increased gluconeogenic potential (PTT assay) (Figure 6J) and elevated fasting blood glucose levels (Figure 6K), when compared to SIRT1-WT expressing mice. Enhanced gluconeogenic potential of SIRT1-mutE2 was further confirmed by measuring glucose output in primary hepatocytes (Figure S6C). We also found elevated expression of PEPCK and G6Pase (Figure 6L), which substantiated the above findings of increased glucose output from livers of SIRT1-mutE2 expressing mice. Furthermore, reduced oil red staining, under two different paradigms of fatty acid feeding and SIRT1 expression, along with elevated expression of genes involved in fatty acid oxidation indicated altered fat metabolism in the absence of SIRT1 glycosylation (Figures S6D-S6F and 6L). SIRT1-mutE2 mice also had decreased levels of lipogenic and fatty acid transporter gene expression even upon 6 hours of re-feeding (Figure S6G). Interestingly, we found elevated expression of inflammatory genes in the liver of SIRT1-mutE2 overexpressing mice (Figure S6H). Highlighting gross perturbation at the level of metabolic signaling, ectopic expression of SIRT1-mutE2 resulted in heightened pAMPK and dampened pAKT (Figure 6M). Taken together, these results demonstrate that altered molecular interactions of SIRT1-mutE2 with PGC1α-PPARα/FOXO and AKT caused physiological defects in the liver.

Since glycosylation was induced under a fed state and led to cytosolic export of SIRT1 followed by degradation, we were curious to check if this modification had any effect on fasted-to-refed transitions. As shown in Figures 6N-6P, SIRT1^co/co^ mice rescued with SIRT1-WT had decreased FOXO1 and pAMPK levels upon re-feeding and a concomitant increase in pAKT. However, putting back SIRT1-mutE2 into SIRT1^co/co^ mice displayed no change in FOXO1, pAKT or pAMPK levels between the starved and the refed states (Figures 6N-6P). Additionally, gluconeogenic and beta-oxidation genes were not only elevated in the SIRT1-mutE2 mice livers, but also did not decrease upon re-feeding when compared to SIRT1-WT (Figure S6I). Together, these results establish that, loss/abrogation of SIRT1 glycosylation leads to continued expression of fasting genes and elevated blood glucose levels, resulting in a failure to transit to a refed state, even upon nutrient availability.

## Discussion

Temporal control of protein-protein interactions is vital for mediating physiological transitions, specifically in the context of dynamic toggling between fed-fast-refed states. Here, we have discovered a nutrient dependent control of SIRT1-functions, which is manifested through a temporal switch in its interactions to regulate hepatic glucose and fat metabolism. In the absence of which, catabolic to anabolic transitions are perturbed leading to liver dysfunctions.

We have now found a regulatory modification on SIRT1, i.e. glycosylation, which determines the interactions of SIRT1 with its target transcription factors/co-regulators. Several studies have identified positive and negative regulators of SIRT1, including proteins like DBC1 (Kim et al., 2008; Krzysiak et al., 2018) and modifications such as sumoylation and phosphorylation (Revollo and Li, 2013). However, mechanistic insights into the spatial and temporal regulation of substrate selectivity of SIRT1, during physiological transitions such as fed-fast-refed cycles, remain largely unknown. In this regard, our results demonstrate that SIRT1 glycosylation during the initial phases of refed state reduces its interaction with nuclear transcription factors/co-regulators (PPARα, PGC1α, FOXO1 and SREBP1), results in cytoplasmic export and enhanced interaction with AKT, a key component of insulin signaling.

The N-terminal specificity domain in SIRT1, which we have recently identified to be encoded by exon-2 (Deota et al., 2017), is part of the IDR. Modifications on IDRs have been thought to further contribute to protein-protein interactions (Babu, 2016). In this context, the results presented here, along with our recent report (Deota et al., 2017), and indicates that metabolite driven modification of exon-2 domain acts to modulate the functions of this master regulator of metabolic gene transcription. In a recent paper, Krzysiak et al demonstrated that binding of DBC1-/PACS2 to the N-terminal IDR in SIRT1, in response to insulin signaling, is involved in its inhibition and thus attenuation of starvation responsive gene transcription. However, if/how a fed response leads to gating or attenuation of SIRT1 functions by altering its interactions and stability has not been addressed thus far. In this context, our results establish that glycosylation of SIRT1 in the exon-2 domain is an important determinant of interaction specificities with transcription factors and signaling molecules in a time dependent manner. Moreover, our findings highlight a direct interplay between glucose inputs and SIRT1 regulation, which seems to be upstream and is required to sensitize insulin signaling itself. Importantly, mutating glycosylation sites on SIRT1 exon-2 domain led to stabilization of its interactions with transcription factors and reduced binding to AKT, which together affected both gene transcription and insulin signaling.

Recent literature has emphasized the importance of fed-fast cycles in addition to drawing attention to restricted feeding regimes (Chaix et al., 2019; Chaix et al., 2014; Hatori et al., 2012; Satoh et al., 2006). Some of these reports have clearly highlighted the fact that aberrant fed-fast cycles lead to metabolic diseases and specifically compromise liver functions (Kinouchi et al., 2018; Longo and Panda, 2016; Sulli et al., 2018). While nutrient and endocrine inputs have been implicated, if/how such inputs encode temporal molecular switches in enabling such transitions are poorly understood. In this context, we establish that abrogating SIRT1 glycosylation results in enhanced fatty acid oxidation and gluconeogenesis, which contribute to perturbed glucose homeostasis both under fasted and refed conditions. Moreover, in addition to failure to transit to a refed state liver expressing non-glycosylatable SIRT1 displayed signs of hyper-inflammation.

Lastly, our study brings to the fore the importance of SIRT1 oscillation both in terms of interactions and levels during fed-fast cycles. Given that there have been many attempts to create gain-of-function models of SIRT1 (Banks et al., 2008; Bordone et al., 2007; Herranz and Serrano, 2010; Hubbard and Sinclair, 2014; Kawashima et al., 2011; Milne et al., 2007), our study points out that uncontrolled activation of SIRT1, as in the case of hypoglycosylation, can be as detrimental to physiological homeostasis. However, we have also seen that glycosylation is positively correlated with reduced SIRT1 levels during aging and in obesity. This suggests that, although glycosylation is necessary to restrict SIRT1 activity upon refeeding, excess glycosylation might lead to a loss of SIRT1 function. We propose that developing interventions, which regulate SIRT1 glycosylation and hence bias its interactions or maintain its homeostatic expression will have therapeutic value.

## ACKNOWLEDGEMENTS

We acknowledge the following funding sources: DAE-TIFR (Govt. of India) (Grant Number 12P-0122) to UKS. We acknowledge IISER and TIFR animal facilities along with Dr. Kalidas Kohale, Dr. Shital Suryavanshi, Dr. Sachin Atole, Dr. Sagar Tarate and Ms. Ritika Gupta for the help with animal experiments.

## AUTHOR CONTRIBUTIONS

Conceptualization, U.K.S.; T.C; S.B; Investigation, U.K.S.; T.C; B.M; S.B; A.C; H.P.B; Writing – Original Draft, U.K.S.; T.C; B.M; S.B; Funding acquisition, U.K.S.; Supervision, U.K.S.

## Material and Methods

### Animals

C57BL/6 and SIRT1^lox/lox^ mice were housed under standard animal house conditions. 3-4 month old and 20-22 month old male mice were considered as young and aged cohorts, respectively. The procedures and the project were approved and were in accordance with the institute animal ethics committee (IAEC) guidelines. Mice were given injections intravenous (i.v.), via the tail vein, of Ad-SIRT1-WT, AdSIRT1-mutE2 or AdCMV. For C57BL/6 mice, i.v. was performed on Day1, followed by fasting blood glucose estimation on Day 4. Time Restricted Feeding (TRF) paradigm was followed for 15 days, in which feed was added at night-break i.e. 7pm, and removed at day-break i.e. 7am, after which experiments were performed as indicated in Figure S6A. Liver tissues were harvested at 1am (6h post feeding) for gene expression and western blot analysis. For SIRT1^lox/lox^ mice, adenovirus overexpressing Cre-recombinase was injected i.v. on Day 1, followed by Ad-SIRT1-WT, Ad-SIRT1-mutE2 or Ad-CMV administration on Day 5. Post injection, experiments were performed as indicated in Figure S6B. For this cohort, liver tissues were harvested either after 24h of starvation or 6h post feeding (6h-RF). Dissected tissues were either snap frozen in liquid nitrogen or processed for protein lysates and total RNA extraction.

### Plasmids and Constructs

pAd-Track FLAG-SIRT1 was a gift from Pere Puigserver (Addgene plasmid # 8438). pAdtrack CMV plasmid was a gift from Bert Vogelstein (Addgene plasmid # 16405). SIRT1-mutE2 (T^160^/S^161^-A) construct generation was outsourced to Genscript Inc. pAd-Track FLAG-SIRT1- mutE2 was generated by sub-cloning -mutE2 from pcDNA3.1 Myc-SIRT1-mutE2 into pAd- Track FLAG-SIRT1. pAd-Track FLAG-SIRT1-ΔE2 was generated by sub-cloning from pcDNA3.1 Myc-SIRT1-ΔE2 into pAd-Track FLAG-SIRT1 (Deota et al., 2017). pLKO-puro FLAG SREBP1 was a gift from David Sabatini (Addgene plasmid # 32017). PPRE-X3-TK-luc was a gift from Bruce Spiegelman (Addgene plasmid # 1015). FAS promoter luciferase was a gift from Bruce Spiegelman (Addgene plasmid # 8890). pcDNA4 myc PGC-1 alpha was a gift from Toren Finkel (Addgene plasmid # 10974). pSG5 PPAR alpha was a gift from Bruce Spiegelman (Addgene plasmid # 22751). FHRE-Luc was a gift from Michael Greenberg (Addgene plasmid # 1789). HA-OGT was a gift from Prof. Gerald W. Hart, Johns Hopkins.

### Cell culture and transfection

HEK-293T, HeLa and HepG2 cells were cultured in DMEM (Sigma) with 10% fetal bovine serum (Gibco) at 37°C under 5% CO2. Indicated plasmids were transfected with Lipofectamine-2000 (Invitrogen) (for 36-42 hours), as per manufacturer’s instructions. Cells were incubated with either low glucose medium (LGM, 5mM glucose) or high glucose medium (HGM, 25mM glucose) for 12h, with fresh HGM supplemented 3h prior to cell collection. Adenoviral particles were used for transducing cell lines after 12h of plating; cells were collected 36-40 hours post transduction after respective treatments. Primary hepatocyte cultures were transduced with adenovirus after 12h of plating and collected at 72h post plating after respective treatments.

### Cell Culture Treatments

All treatments were given in 10% FBS containing media unless otherwise specified. For OGA inhibition experiments, PUGNAc was used at a concentration of 100µM for 12-16h in HGM. For OGT inhibition, cells were treated with 100µM BZX for 16h under low/high glucose conditions, as indicated in individual figures. For insulin stimulation assays, cells were kept in 5% FBS containing DMEM medium overnight followed by 1h in serum free EBSS prior to addition of 100nM insulin in high glucose conditions for time points indicated in respective figures. MG132 (20µM) was used for inhibition of proteasome for either 12-16h or at time points indicated in the figures. Cyclohexamide [CHX (100µg/ml)] was added to cell at time points indicated in individual figures for inhibition of protein translation. CRM-1 mediated nuclear export was inhibited using Leptomycin B (LMB) at a concentration of 10ng/ml for 12-16h. Adenoviral SIRT1-WT and mutE2 overexpression in hepatocytes and HeLa cells were done for 24-48h before cell collection.

### Adenoviral preparation

Adenovirus FLAG-SIRT1 was prepared as per the protocol described by Luo et al. (Luo et al., 2007). Cells were collected post expression of GFP and were lysed using hypotonic buffer [HEPES (100mM, pH 7.5), MgCl2 (1.5mM), KCl (10mM), DTT (0.5mM)]. Supernatant was collected and used for transduction in different cell types. SIRT1 expression was confirmed by western blot and qPCR.

### Free Fatty Acid Conjugation and treatment

4.6 g of fat-free BSA was dissolved in 20 ml of 150mM NaCl and divided into two tubes of equal volume. To the first tube, 10ml of 150mM NaCl was added (pH adjusted to 7.4), filter sterilized and stored at −20°C. 61.2 mg of Sodium Palmitate was added to 10 ml of 150mM NaCl and dissolved by heating at 70°C. 8ml of this solution was added to the second tube of BSA solution and kept at 37°C for 1h to make a stock of 10mM. After dissolution, the pH was adjusted to 7.4 using NaOH and volume made upto 20 ml and stored at −20°C. For experiments, 500µM of Palmitate-BSA or BSA was either added to hepatocytes for 12h followed by SIRT1-WT/-mutE2 expression for 24h, or added to SIRT1-WT/-mutE2 over-expressing cells (12h) for 24h, after which oil red staining was done for both the experimental paradigms.

### Lysate preparation

Tissue and cell lysates were prepared in RIPA buffer (50mM Tris pH 8.0, 150mM NaCl, 0.1% SDS, 0.5% Sodium deoxycholate, 1% Triton × 100, 0.1% SDS, 1 mM PMSF, Protease inhibitor cocktail) and debris was pelleted at 12,000 rpm (4°C/10-minutes). Protein concentration was determined as per manufacturer’s instructions and the samples were boiled in SDS gel loading buffer.

### Immunoprecipitation

Lysates in TNN [50mM Tris (pH 7.5), 150mM NaCl, 0.9% NP-40] buffer were pelleted by centrifugation (12K-rpm/15-minutes/4°C), protein estimated from supernatant, pre-cleared using Protein A/G beads for 1h at 4°C and then incubated with indicated antibodies (or Ab-conjugated-beads) overnight. Immune complexes were pulled down either directly or with protein-A/G beads, washed with TNN and eluted in SDS gel loading buffer.

### Nucleo-cytoplasmic fractionation

Cells were lysed in hypotonic lysis buffer (10 mM HEPES pH 7.5, 1.5 mM MgCl2, 10 mM KCl, 0.5 mM DTT, 1 mM PMSF, Protease inhibitor cocktail) and nuclei were pelleted at 2000 rpm at 4°C. The supernatant was re-centrifuged at 12,000 rpm for 15 min and the final supernatant collected as cytosolic extract. The nuclei were washed with ice-cold hypotonic buffer and lysed in RIPA buffer (50mM Tris pH 8.0, 150mM NaCl, 0.1% SDS, 0.5% Sodium deoxycholate, 1% Triton × 100, 0.1% SDS, 1 mM PMSF, Protease inhibitor cocktail) to get the nuclear extract. Nuclear extracts were either, boiled in SDS gel loading buffer and loaded onto SDS-PAGE or used for immunoprecipitation with indicated antibodies.

### Oil Red O (ORO) staining

Primary hepatocytes (6.5×10^4^) were plated onto 24 well collagen coated plates. After respective treatments, cells were rinsed once with PBS and fixed in chilled 4% PFA for 30 minutes. The cells were rinsed once again in PBS and freshly prepared and filtered ORO solution (40% in distilled water from a 0.5% stock solution in isopropanol) was added and kept for 15 minutes. The cells were washed in distilled water and imaged on EVOS FLc microscope from Life technologies Inc. For quantification, ORO was extracted in 100% isopropanol for 15 minutes with shaking. The solution was mixed with pipetting and distributed in 3 wells as triplicates in a 96-well plate and absorbance was read at 492nm. 100% isopropanol was used as background control. The readings were normalized to total protein amount obtained from the cells lysed using RIPA buffer.

### Immunoprecipitation/purification of FLAG-mSIRT1-WT/mut-E2 for activity assays

HEK293T cells were transfected in 150 mm^2^ culture dishes with either FLAG-SIRT1-WT or FLAG-SIRT1-mutE2 constructs. After indicated treatments, cells were washed and lysed in TNN buffer and Immunoprecipitation was done using Anti-FLAG-M2 beads. Elution was performed with 100 uL of 10 mM Tris pH 8.0, 150 mM NaCl, 150 µg/mL FLAG peptide on a thermomixer (16° C, 3h, 1000 rpm). To determine relative expression of the eluted fusion proteins, SDS-PAGE western blotting was done using diluted eluate, anti-Flag antibody and anti-mouse-HRP secondary antibody, chemiluminescent detection and imaging on GE Amersham Imager 600. Expression ratio of SIRT1-FL/-mutE2 was estimated via densitometry in ImageJ (NIH, USA), and loads in subsequent assays were adjusted proportionally. Normalized SIRT1 amount was used for activity assays using calf thymus histones (Roche), as indicated below.

Mouse FLAG-SIRT1 (FL or mutE2; balanced by FLAG-SIRT1 Western expression) was incubated with 0.1mg/ml of Histones in the presence of 0.2 mM NAD^+^, 25mM Tris pH-8.0, 137mM NaCl, 2.7mm KCl and 1mM MgCl_2_ at 37°C for either 30 or 60 minutes. Reaction was quenched at indicated time points with the addition of SDS gel loading buffer and heating at 95°C for 10 min. Samples were loaded onto SDS-PAGE and western blotting was done for indicated proteins.

### Luciferase assays

Cells were transfected with the indicated Promoter luciferase constructs along with β-Gal plasmid. After 24h of transfection, cells were harvested and lysed in Passive Lysis buffer. β- galactosidase assay was performed using the ONPG method. Luciferase assay was performed using the Luciferase Assay system. The luciferase counts were measured in TriStar LB941 (Berthold technologies), and normalized to the β-galactosidase activity and represented as fold change.

### Immunoblotting

Equal amounts of protein were run on a SDS-PAGE and transferred to polyvinylidene difluoride (PVDF) membranes. Following blocking with 5%-milk, they were probed with appropriate primary and secondary antibodies, according to standard procedures. Chemiluminescence detection kit was used to visualize the bands using X-ray films (Fiji films) or GE Amersham Imager 600. Actin and Tubulin levels were used as loading controls, as indicated in respective figures. Band intensities were quantified using ImageJ.

### RNA extraction, RT-PCR and qPCR analysis

Total RNA was extracted from cells and tissues using Trizol reagent. 1 µg of RNA was reverse transcribed using random hexamers and SuperScript-IV RT kit. Quantitative PCR (qPCR) was performed using KAPA SYBR® FAST Universal 2X qPCR Master Mix, and Roche Light Cycler 480 II and LC 96 instruments. RNA isolation, cDNA synthesis and real-time PCR were carried out as per manufacturer’s instructions. The primer pairs used for the assays are tabulated in Supplementary Table S1 in the supplementary information. Levels of 18S were used for normalization.

### Gluconeogenesis assay

6.5×10^4^ hepatocytes from control mice were plated in a 24-well plate. 24h post plating, SIRT1- WT and –mutE2 overexpressing adenovirus was added to the cells. 36h post plating, glucose free media supplemented with Glutamine (2mM) and either 2mM or 4mM pyruvate was added to the cells and incubated for 12h. 500µl of the culture supernatant was collected and glucose levels were measured as per manufacturer’s protocol. In brief, 25µl of the culture supernatant was diluted in assay buffer and glucose levels were estimated and normalized to the protein amounts from total cell lysates.

### Hepatocyte Isolation

Hepatocytes were isolated from male wild-type, 8-12 wk old C57BL/6 mice by collagenase perfusion through the inferior venacava/portal vein as per method described in Maniyadath et al. (Maniyadath et al., 2019). Hepatocytes were counted and plated for respective experiments and treatments were given as indicated in each figure.

### Immunofluorescence

HeLa cells were rinsed thrice with PBS and fixed in ice cold 4% PFA for 10 minutes. After fixation, the cells were washed thrice with PBS, followed by permeabilization with 0.1% Triton X-100 for 10 minutes. After washing with PBS, blocking for 1h was done in 5% BSA at room temperature. Cells were then incubated overnight at 4°C, with indicated primary antibody, followed by washing with PBS and addition of anti-rabbit Alexa Fluor 594 for 45 minutes at RT. Coverslips were then mounted on a slide and imaged at 60X using FluoView® FV1200 Laser Scanning Confocal Microscope from Olympus Life Science.

### Pyruvate and Insulin Tolerance Test (PTT and ITT)

Mice were fasted for 6h, after which body weight and blood glucose levels were measured. 2g/Kg b.wt of pyruvate in PBS (for PTT) or Insulin (0.75 IU/kg b.wt, HumulinR, Lilly) (for ITT) was injected into each mouse intraperitoneally. The blood glucose level was monitored with a glucose meter at indicated intervals during a 2h time course.

### Glucose Tolerance Test (GTT)

Mice were fasted for 12-16h, after which body weight and blood glucose levels were measured. 2g/Kg b.wt of glucose was injected into each mouse intraperitoneally. The blood glucose level was measured with a glucose meter at indicated intervals during a 2h time course.

### Measurement of Palmitate induced fatty acid oxidation (FAO)

FAO was measured as Palmitate induced Oxygen consumption rate (OCR) using Seahorse XFe24 Extracellular flux analyzer, in cultured primary hepatocytes (1.8×10^4^ cells/well). Cells transduced with control, SIRT1-WT and -mutE2 adenovirus were treated with substrate limiting media [DMEM containing 0.5mM glucose, 0.5mM Carnitine, 1mM glutamine and supplemented with 1% FBS] for 24h. Prior to the assay, cells were treated with FAO assay medium (KHB containing 2.5mM glucose, 0.5mM Carnitine and 5mM HEPES) for 45 minutes. To measure beta-oxidation independent OCR, cells were pre-treated for 15 minutes with 60µM of Etomoxir (CPT1 inhibitor) and was retained during the assay period. Cells were incubated for 45 minutes at 37°C in a non-CO2 incubator as per manufacturer’s instructions. Just prior to the assay, 200µM of Palmitate conjugated with BSA was added as substrate for beta-oxidation. OCR (pmol/min) was measured in 3 cycles each for basal rate and following injections of 2µM oligomycin and 1µM Rotenone. Readings from 3 technical replicates were normalized to blank wells in the plate and to cell numbers in respective wells, and the assays were repeated at least thrice. Results from all the experimental repeats were used for calculating Palmitate induced basal OCR and ATP production rate.

### Measurement of extracellular acidification rate (ECAR)

HeLa cells (1.8×10^4^ cells/wells) plated in HG-DMEM were transduced with control, SIRT1-WT and -mutE2 adenovirus in the Seahorse XFe24 Extracellular flux analyzer plates. Post 36h, cells were treated with NG-DMEM supplemented with 2mM Glutamine and 2mM Sodium pyruvate, for 4h. Prior to the assay, cells were washed and incubated with Glycostress assay medium (XF Basal media supplemented with 2mM Glutamine) for 45 minutes at 37°C in a non-CO2 incubator. ECAR (mpH/min) was measured in multiple cycles for basal rate and following 10mM D-Glucose injection. Acute acidification rate in response to insulin was measured by injecting 100nM Insulin and non-glycolytic ECAR was determined by injecting 100mM 2- Deoxy-Glucose. Readings from 3 technical replicates were normalized to blank wells in the plate and to cell numbers in respective wells, and the assays were repeated at least twice. Results from all the experimental repeats were collated for computing Relative Basal ECAR and Acute Insulin Response.

### QUANTITATION AND STATISTICAL ANALYSIS

Data are expressed as means ± standard error of means (SEM). Statistical analyses were performed using Microsoft Excel (2013) and Graph Pad Prism (version 6.0). Statistical significance was determined by the Student’s t test. A value of p ≤ 0.05 was considered statistically significant. *p≤ 0.05; **p≤ 0.01; ***p≤ 0.001.

## SUPPLEMENTAL INFORMATION

Supplemental information includes six figures and one table, and can be found with this article.

**Table S1:** qPCR primers used in the study.

**Figure S1: Nutrient dependent regulation of SIRT1 protein stability and glycosylation. Related to Figure 1**. (A-E) Immunoblot for endogenous SIRT1 in (A) HepG2 cells and (B) Primary hepatocytes, treated with 100nM Insulin for indicated time points; (C) 24h starved and 6h-refed liver tissue; (D) Normal chow diet (ND) fed and 3-months High Fat Diet (HFD) fed mice liver tissue and (E) Young (3 months) and old (20 months) mice liver tissue. (F) Immunoblot for glycosylated SIRT1 enriched in sWGA pull down from HEK293T cells. (G-H) FLAG-M2 immunoprecipitates from HEK293T cells treated with 25mM 2-deoxy glucose (2- DG), expressing FLAG-SIRT1-WT and immunoblotted for (G) glycosylated SIRT1 and (H) interaction with HA-OGT. (I) Immunoblot for glycosylated SIRT1 of FLAG-M2 immunoprecipitates from HEK293T cells expressing FLAG-SIRT1-WT treated with 100µM BZX.

**Figure S2: Identification and characterization of SIRT1 Exon-2 glycosylation sites. Related to Figure 2**. (A) *In silico* prediction of glycosylation sites in Exon-2 domain of SIRT1 using YingOYang 1.2 software. (B) Schematic representation of cloning strategy for generating Thr^160^/Ser^161^ to Ala mutants. (C) Immunoblot for interactions of SIRT1-WT, ΔE2 and mutE2 with OGT in HA immunoprecipitates from HEK293T cells expressing FLAG-SIRT1-WT, -ΔE2, –mutE2 and HA-OGT. (D) Immunoblot for glycosylated SIRT1 in FLAG-M2 immunoprecipitates from HEK293T cells expressing FLAG-SIRT1-WT and -ΔE2. (E-F) Comparison of SIRT1 activity from FLAG-M2 immunoprecipitates in HEK293T cells overexpressing FLAG-SIRT1-WT (± PUGNAc) and –mutE2, immunobloted for (E) Endogenous SIRT1 and (F) Acetylated H3K9.

**Figure S3: Glycosylation dependent regulation of mitochondrial fatty acid oxidation, insulin signaling and glycolysis. Related to Figure 3**. (A-B) Palmitate-induced oxygen consumption rate (OCR-FAO) in primary hepatocytes expressing FLAG-SIRT1-WT/-mutE2 and quantified for (A) Basal respiration (n=3, N=2) and (B) ATP production rate (n=3, N=2). (C) Immunoblot for insulin signaling kinetics in HeLa cells expressing either FLAG-SIRT1-WT or - mutE2 and treated with 100nM Insulin for indicated time points. (D-E) Extracellular Acidification Rate (ECAR) in HeLa cells expressing either FLAG-SIRT1-WT or -mutE2 under (D) Basal conditions (n=3, N=2) and (E) in response to administration of 100nM insulin (n=3, N=2). Data are represented as mean ± SEM and analyzed by the Student’s t test. A value of P ≤ 0.05 was considered statistically significant. #,*P≤ 0.05; ##, **P≤ 0.01; ***P≤ 0.001. * and # denotes a comparison w.r.t Control and SIRT1-WT, respectively.

**Figure S4: Nucleo-cytoplasmic localization of glycosylated SIRT1. Related to Figure 4**. (A) Immunoblot for glycosylated SIRT1 in different cellular fractions from either sWGA purified or FLAG-M2 immunoprecipitates. (B) Dosage standardization of Leptomycin B treatment (12h) in HeLa cells. Scale bars represent 20µm. (C) SIRT1 immunoprecipitates from nuclear fraction of HeLa cells treated with 100µM PUGNAc and 10ng/ml LMB for 12h, immunoblotted for glycosylated SIRT1. T=Total cell lysate; C=Cytoplasmic fraction; N= Nuclear fraction.

**Figure S5: SIRT1-WT and –mutE2 protein stability in response to inhibitor treatments. Related to Figure 5**. (A) Immunoblot and quantification of endogenous SIRT1 in HeLa cells treated with ± BZX (100µM, 16h) and CHX (20µM) for indicated time points. (B) Immunoblot and quantification of FLAG-SIRT1-WT and –mutE2 in HeLa cells treated with PUGNAc (100µM) and CHX (20µM) for indicated time points. (C) Immunoblot for endogenous SIRT1 in HeLa cells treated with LMB (10ng/ml, 12h), CHX (20µM, indicated time points) and ± PUGNAc (100µM, 16h). (D) Immunoblot for SIRT1 in HeLa cells overexpressing FLAG- SIRT1-WT/–mutE2 treated with and PUGNAc (100µM, 16h), LMB (10ng/ml, 12h) and CHX (20µM) for indicated time points.

**Figure S6: Effect of SIRT1-WT and –mutE2 in regulating cellular and organismal physiology. Related to Figure 6**. (A-B) Schematic representation of experimental paradigm used for (A) hepatic overexpression of FLAG-SIRT1-WT and –mutE2 in C57Bl6 mice and (B) hepatic rescue of SIRT1 expression using Ad-SIRT1-WT and –mutE2 in SIRT1^co/co^ mice. (C) Gluconeogenesis assay in primary hepatocytes overexpressing either SIRT1-WT or –mutE2 and treated with 2mM and 4mM pyruvate (n=4, N=2). (D-F) Oil-Red O (ORO) staining experiments in primary hepatocytes (D-E) overexpressing either SIRT1-WT or –mutE2 followed by loading with Free Fatty Acid (FFA, 500µM) (D) representative images for SIRT1-WT and –mutE2 overexpression and ORO staining and (E) Colorimetric quantification of ORO stain (n=3, N=2); (F) Colorimetric quantification of ORO stain in hepatocytes loaded with Free Fatty Acid (FFA, 500µM) followed by overexpression of either SIRT1-WT or –mutE2 (n=3, N=2). (G-H) Gene expression analysis in C57Bl6 mice livers overexpressing SIRT1-WT and –mutE2 for (G) lipogenic and fatty acid transporter genes (n=3, N=3), and (H) inflammatory genes (n=3, N=3). (I) Gluconeogenic and fatty acid oxidation gene expression analysis from 24h starved and 6h- refed livers of SIRT1^co/co^ mice rescued with Ad-SIRT1-WT and –mutE2 (n=3). Data are represented as mean ± SEM and analyzed by the Student’s t test. A value of P ≤ 0.05 was considered statistically significant., *P≤ 0.05; **P≤ 0.01; ***P≤ 0.001.

