## Supplemental information and Figures for "Temporal gating of SIRT1 functions by O-GlcNAcylation prevents hyperglycemia and enables physiological transitions in liver"

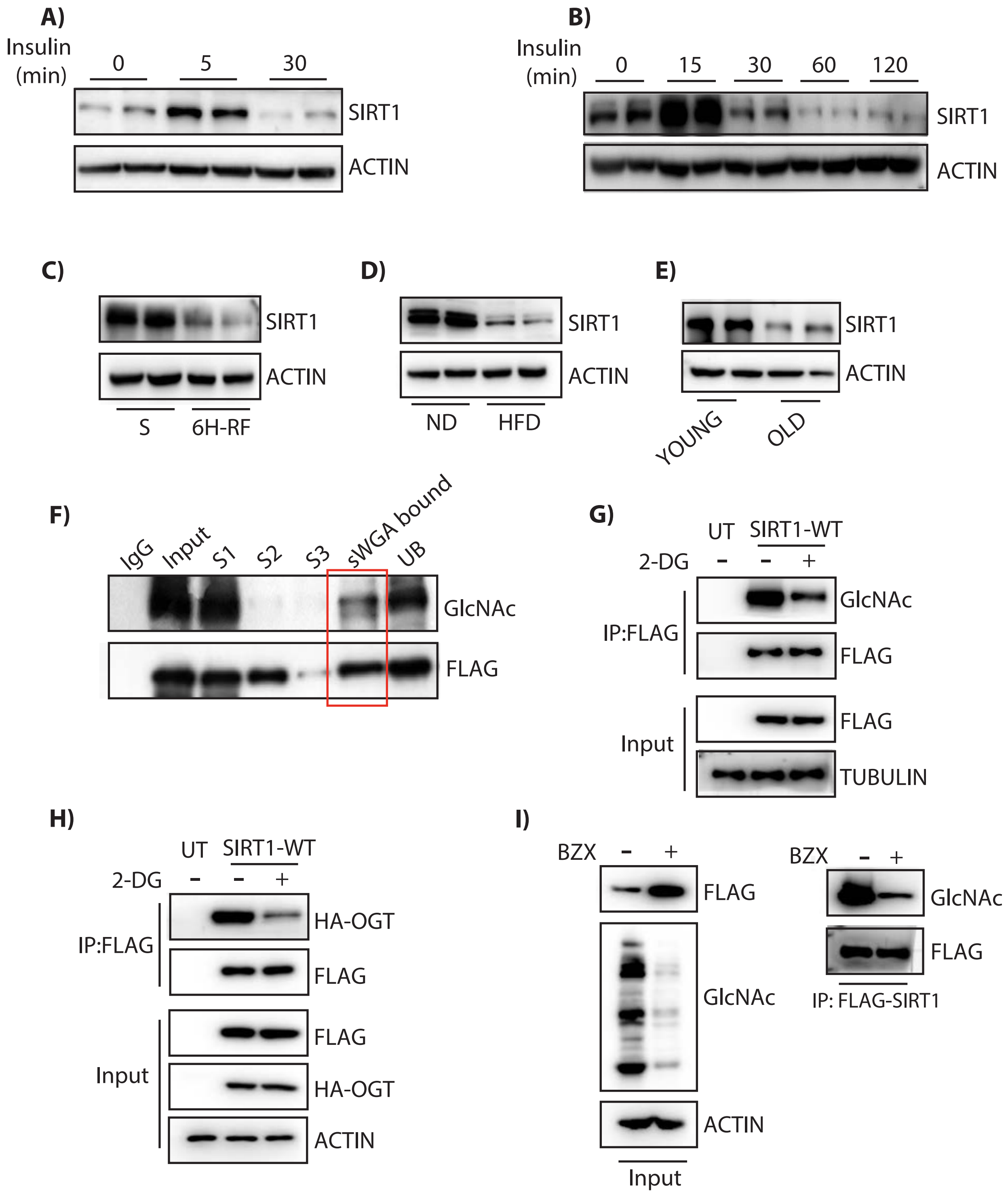


**Figure S1. Related to Figure 1**

A)

| Rank | Position | AA | Potential O-GlcNAc |
|---|---|---|---|
| 1 | 161 | S | 0.6284 |
| 2 | 658 | S | 0.5607 |
| 3 | 257 | S | 0.5578 |
| 4 | 160 | T | 0.5497 |
| 5 | 657 | S | 0.5078 |
| 6 | 446 | S | 0.4844 |
| 7 | 596 | S | 0.4528 |
| 8 | 737 | S | 0.4202 |

B)

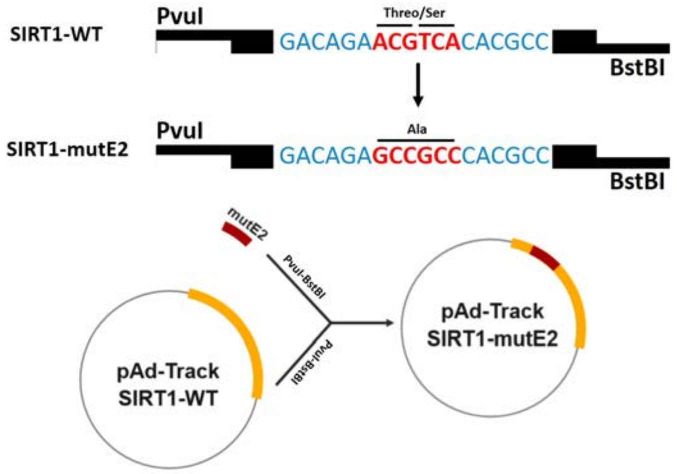


C)

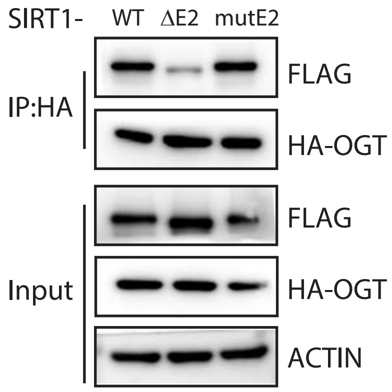


D)

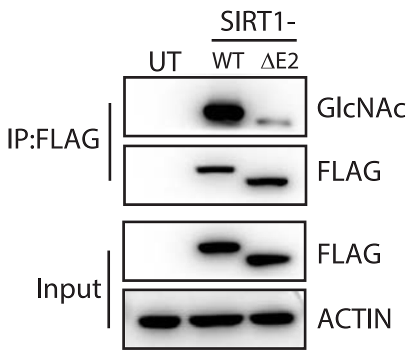


E)

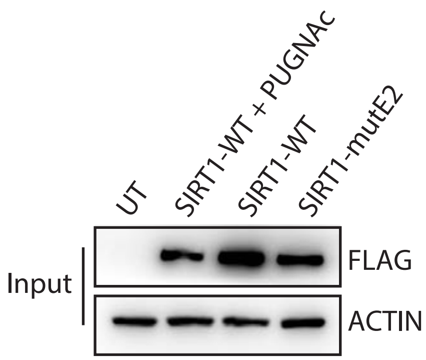


F)

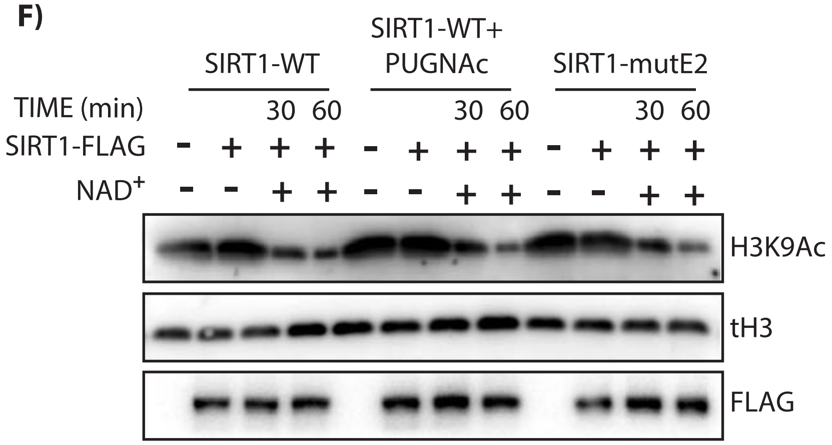


**Figure S2. Related to Figure 2**

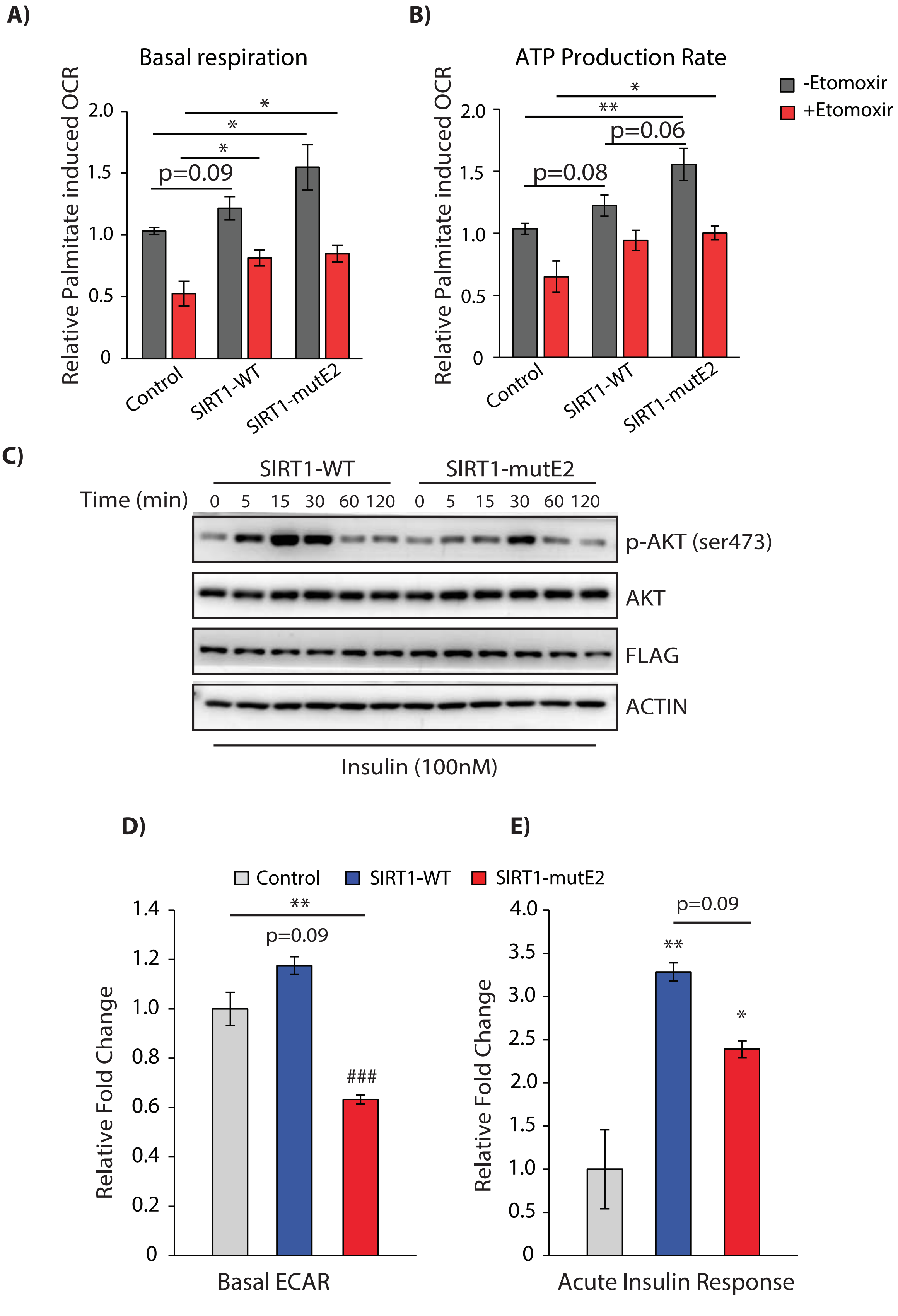


**Figure S3. Related to Figure 3**

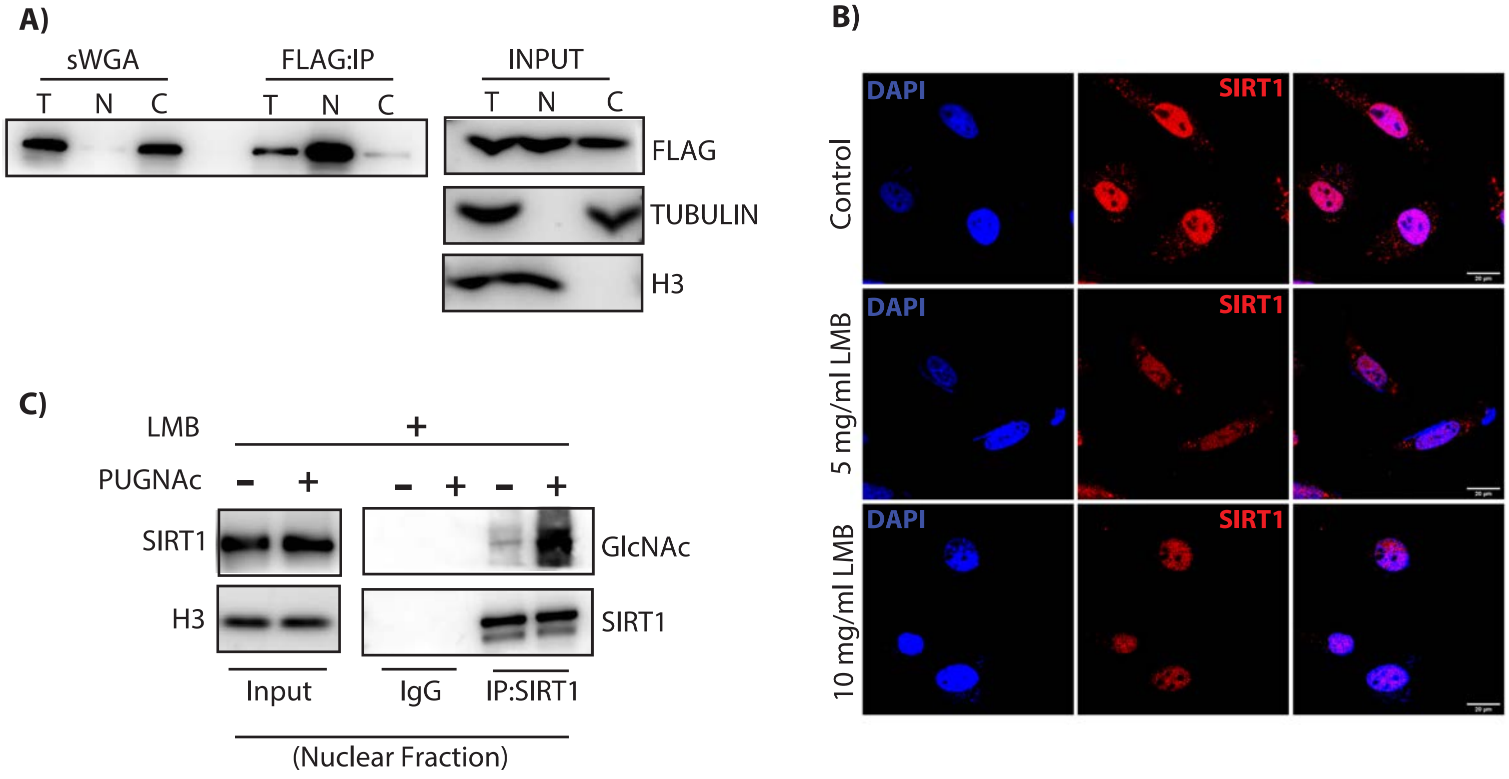


**Figure S4. Related to Figure 4**

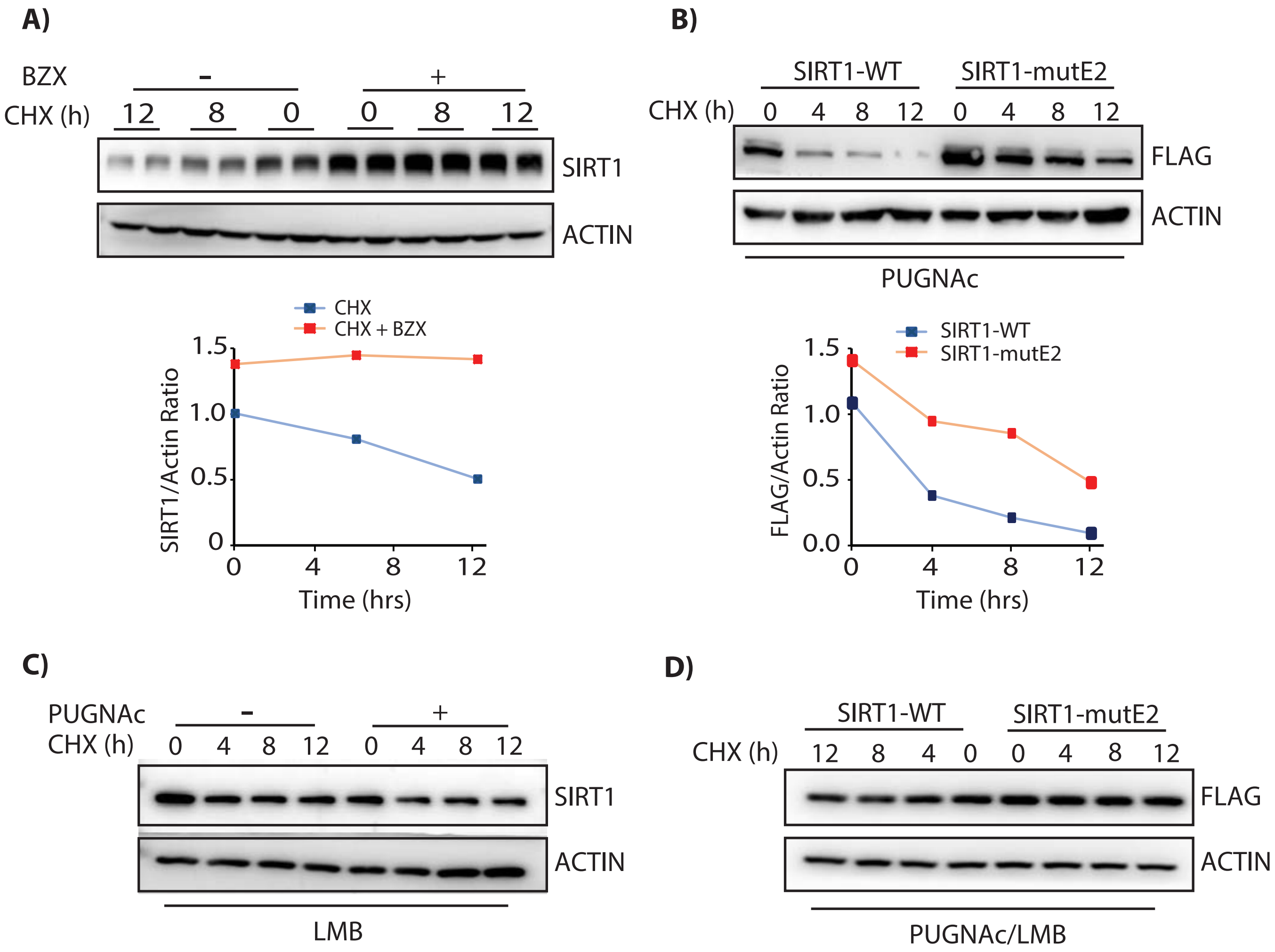


**Figure S5. Related to Figure 5**

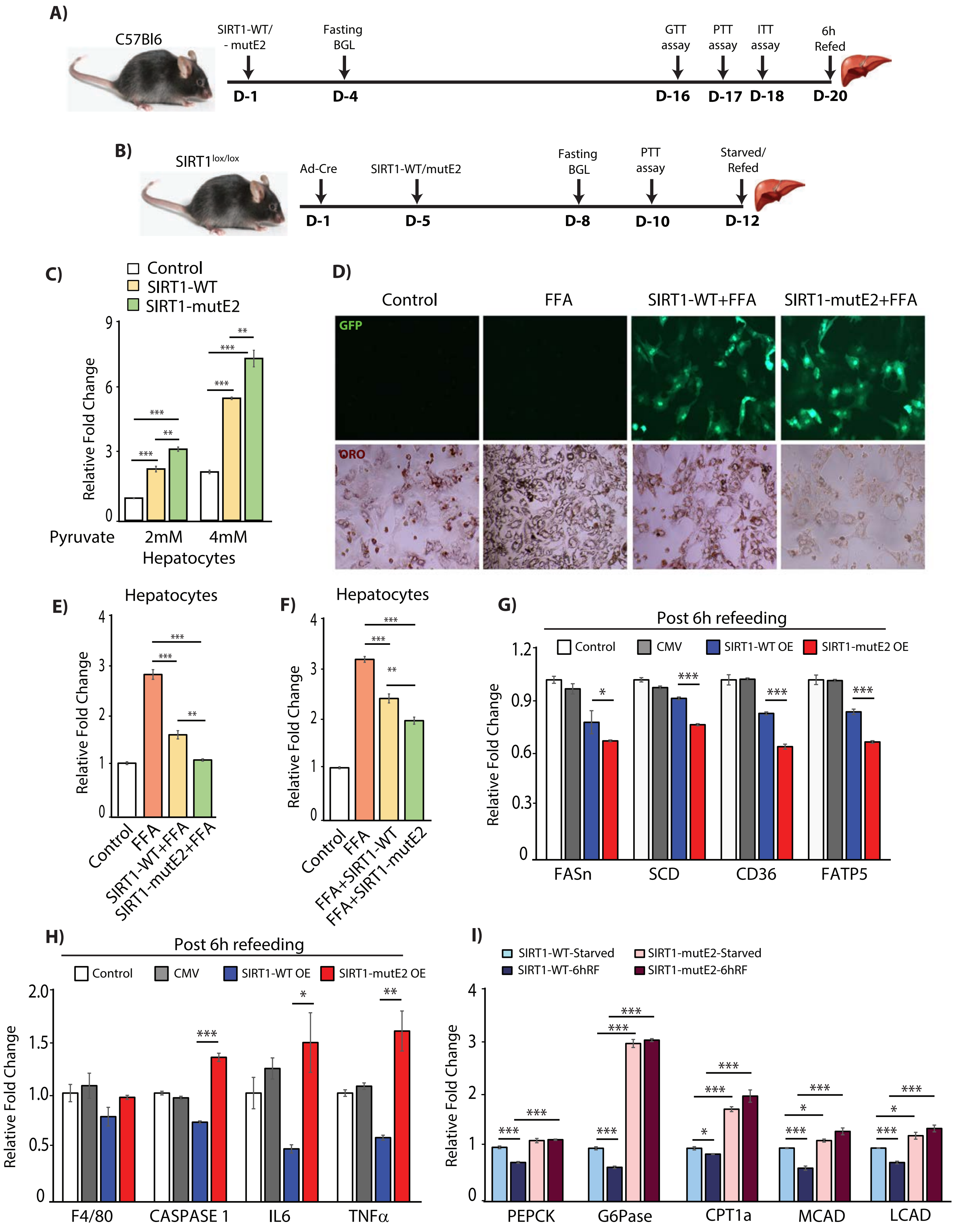


**Figure S6. Related to Figure 6**
