## Supplemental Table 1 for "Temporal gating of SIRT1 functions by O-GlcNAcylation prevents hyperglycemia and enables physiological transitions in liver"

| **GENES** | **PRIMER SEQUENCES** |
| --- | --- |
| SIRT1 | FP: 5’- GTAACCCTGTAAAGCTTTCAG -3’  RP: 5’- CAGAAGAGTCTTGTGGTACAG -3’ |
| PGC1α | FP: 5’- GTGGATGAAGACGGATTGCC -3’  RP: 5’- GCTGAGTGTTGGCTGGTGCC -3’ |
| SREBP1c | FP: 5’- CTGCTGGACCACAGAAAGGT -3’  RP: 5’- GTAGAGGCTAAGCTGTCCCG -3’ |
| FAS | FP: 5’- TTGCTGGCACTACAGAATGC -3’  RP: 5’- ACTCCCTGAATCATCAAAGG -3’ |
| LCAD | FP: 5’- TCTTTTCCTCGGAGCATGACA-3’  RP: 5’- GACCTCTCTACTCACTTCTGA -3’ |
| CPT1β | FP: 5’-TCGTCACCTCTTCTGCCTTT-3’  RP: 5’-ACACACCATAGCCGTCATCA-3’ |
| PEPCK | FP: 5’-TTCGGCAAATACCTGGCCCACT-3’  RP: 5’-CCCCAGGCCTTTCAGGTTCA-3’ |
| MCAD | FP: 5’-TTGAGTTCACCGAACAGCAG-3’  RP: 5’- ATCCGCTGCACAGATCCAAA-3’ |
| FASn | FP: 5’- TTGCTGGCACTACAGAATGC -3’  RP: 5’- ACTCCCTGAATCATCAAAGG -3’ |
| ACC | FP: 5’- AAGGCTATGTGAAGGATGTGG -3’  RP: 5’- CTGTCTGAAGAGGTTAGGGAAG-3’ |
| SCD1 | FP: 5’- CTGACCTGAAAGCCGAGAAG -3’  RP: 5’- AGAAGGTGCTAACGAACAGG -3’ |
| G6Pase | FP: 5’- GAGGAAGGAATGAACATTCT -3’  RP: 5’- TGGGCTTGCTCTTCTGTATC -3’ |
| CPT1a | FP: 5’-GCAGTCGACTCACCTTTCCT-3’  RP: 5’- ATTTCTCAAAGTCAAACAGTTCCA-3’ |
| CD36 | FP: 5’- GCGACATGATTAATGGCACAG-3’  RP: 5’- GATCCGAACACAGCGTAGATAG -3’ |
| FATP5 | FP: 5’- ACGTCCTACCTCTGTACCATAC-3’  RP: 5’- CAAGATCACTGTTACGCCATG -3’ |
| F4/80 | FP: 5’- TGACTCACCTTGTGGTCCTAA-3’  RP: 5’- CTTCCCAGAATCCAGTCTTTCC -3’ |
| Caspase 1 | FP: 5’- AGGCACGGGACCTATGTGAT -3’  RP: 5’- ATTCCTGCCAGGTAGCAGTC -3’ |
| IL-6 | FP: 5’- TCCAGTTGCCTTCTTGGGAC -3’  RP: 5’- GTGTAATTAAGCCTCCGACTTG -3’ |
| TNFα | FP: 5’- CAAAATTCGAGTGACAAGCCTG -3’  RP: 5’- GAGATCCATGCCGTTGGC -3’ |
| 18S | FP: 5’-TTTCGAGGCCCTGTAATTGG-3’  RP: 5’-CCCAAGATCCAACTACGAGC-3’ |

**Table S1:** qPCR primers used in this study. Related to STAR Methods.
